# Interplay between the paradox of enrichment and nutrient cycling in food webs

**DOI:** 10.1101/276592

**Authors:** Pierre Quévreux, Sébastien Barot, Élisa Thébault

## Abstract

Nutrient cycling is fundamental to ecosystem functioning. Despite recent major advances in the understanding of complex food web dynamics, food web models have so far generally ignored nutrient cycling. However, nutrient cycling is expected to strongly impact food web stability and functioning. To make up for this gap, we built an allometric and size structured food web model including nutrient cycling. By releasing mineral nutrients, recycling increases the availability of limiting resources for primary producers and links each trophic level to the bottom of food webs. We found that nutrient cycling can provide a significant part of the total nutrient supply of the food web, leading to a strong enrichment effect that promotes species persistence in nutrient poor ecosystems but leads to a paradox of enrichment at high nutrient inputs. The presence of recycling loops linking each trophic level to the basal resources weakly affects species biomass temporal variability in the food web. Recycling loops tend to slightly dampen the destabilising effect of nutrient enrichment on consumer temporal variability while they have opposite effects for primary producers. By considering nutrient cycling, this new model improves our understanding of the response of food webs to nutrient availability and opens perspectives to better link studies on food web dynamics and ecosystem functioning.

## Introduction

Food web dynamics and functioning have been studied thoroughly through empirical and modelling approaches because food webs are essential to ecosystem functioning. A central issue is to determine the characteristics of food webs that affect their key properties, *e.g.* the number of species composing them, primary production or secondary production. Food chains (*i.e.* linear chains of species or trophic groups interacting through trophic interactions) and food webs (*i.e.* networks of species interacting through trophic interactions) models have been extensively used to tackle these issues. In particular, dynamical models of complex food webs (*i.e.* food webs including numerous interacting species) reveal that size structured food webs (Brose et al., 2006b; Heckmann et al., 2012), allometric scaling of biological rates (Brose et al., 2006b) and adaptive foraging (Kondoh, 2003; Heckmann et al., 2012) promote species coexistence and population stability. However, these models focus on population dynamics and carbon fluxes, forgetting non-living compartments (mineral nutrients and dead organic matter) and nutrient cycling (cyclic fluxes of nutrients through living and non-living compartments). Some studies include mineral nutrients as basal resources for primary producers (Schneider et al., 2016; Wang and Brose, 2017) or detritus as basal resources for bacteria (Boit et al., 2012) or for omnivorous consumers as well (Legagneux et al., 2012), but they never include a complete nutrient cycling.

Nevertheless, the cycling of mineral nutrients such as nitrogen and phosphorus likely tightly interacts with food web stability. Stability can be measured in different ways: resilience (calculated by the leading eigenvalue of the jacobian matrix of the system at equilibrium) represents the ability of the system to return to its equilibrium after a perturbation, resistance measures the degree to which a variable changes after a perturbation and temporal variability (measured for example by the coefficient of variation) represents the variance of population densities over time (McCann, 2000). Several studies highlighted the importance of nutrient cycling processes for ecosystem stability, but with contrasting results (O’Neill, 1976; DeAngelis, 1980; DeAngelis et al., 1989; DeAngelis, 1992; Loreau, 1994; McCann, 2011; Neutel and Thorne, 2014). DeAngelis (1980, 1992) showed that nutrient cycling affects food chain resilience, systems with tighter nutrient cycling (*i.e.* a lower proportion of mineral nutrients is lost from the ecosystem each time they cycle) being less resilient. On the other hand, Loreau (1994) suggested that tighter cycling was associated with greater food chain resistance to perturbations, and McCann (2011) found that food chains with nutrient cycling were less destabilised (*i.e.* more resilient) by nutrient enrichment than food chains without nutrient cycling. Meanwhile, Neutel and Thorne (2014) did not find clear effects of the presence of recycling loops on the resilience of complex soil food webs, some food webs being unaffected by nutrient cycling and others being either destabilised or stabilised. While the study of consequences of nutrient cycling on stability has largely been restricted to resilience of small food web motifs or food chains (but see Neutel and Thorne (2014)), understanding the consequences of nutrient cycling on species dynamics in complex food webs becomes crucial to predict ecosystem response to perturbations. Observed contradictory results on the effects of nutrient cycling might arise from the fact that nutrient cycling can affect food web through different mechanisms, whose importance could also differ between food chain and food web models.

First, the recycled nutrients (*i.e.* excreted nutrients that return to the mineral pool available for primary producers) are added to the external inputs of mineral nutrients and could lead to an enrichment effect (Loreau, 2010). Nutrient availability has contrasting effects on food webs. On one hand, it fuels primary production and increases the energy transfer to consumers, leading to a higher species persistence and sustaining higher trophic levels as supported by models (Abrams, 1993; Binzer et al., 2011) and empirical observations (Yodzis, 1984; Doi, 2012). On the other hand, nutrient overabundance tends to increase the amplitude of population oscillations, which increases the risk of extinction. This characterises the paradox of enrichment (Rosenzweig, 1971; Rip and McCann, 2011) predicted by several food chain and food web models (Roy and Chattopadhyay, 2007; Rall et al., 2008; Hauzy et al., 2013; Gounand et al., 2014; Binzer et al., 2016) and some experiments (Fussmann et al., 2000; Persson et al., 2001). Taken together, this leads to the hypothesis that in nutrient poor ecosystems, nutrient cycling would have a positive effect on food webs, *i.e.* on species persistence and the persistence of higher trophic levels while, in nutrient rich ecosystems, nutrient cycling would destabilise food webs. Thus, the enrichment effect of nutrient cycling may be a major component of its impact on food webs (McCann, 2011). This is particularly meaningful in a context of global nutrient enrichment due to human activities (Vitousek and Reiners, 1975; Smith et al., 1999).

Second, nutrient cycling adds direct feedback loops from all trophic levels to the bottom of food webs. Besides the consequent enrichment effect, these feedback loops may affect biomass dynamics (McCann, 2011; Neutel and Thorne, 2014). Because these feedback loops are positive (Fath and Halnes, 2007; Halnes et al., 2007) they may have a destabilising effect causing an increase in the oscillation amplitude of biomass densities. However, they could have the opposite effect if nutrient cycling leads to asynchronous dynamics of mineral nutrients and primary producers, as found in a food chain model (McCann, 2011). In such case, a decrease in primary producers could be dampened by a simultaneous increase in mineral nutrients availability, thus reducing population oscillations in the food chain (Brown et al., 2004a). Such effects of recycling feedback loops on stability might however be weaker in complex food webs. In complex food webs, recycled nutrient inputs to detritus and mineral nutrient pools result from many feedback loops, which might attenuate the fluctuations of mineral nutrient dynamics and thus limit the stabilising (resp. destabilising) effect of asynchronous (resp. synchronous) fluctuations of mineral nutrients and primary producers.

Third, the effects of nutrient cycling on stability might be modulated by the ways nutrient are recycled. Consumers in food webs directly affect nutrient cycling both through immobilisation of nutrients in their biomass and through egestion and excretion of non-assimilated food (Vanni, 2002). Furthermore, nutrients are excreted as mineral nutrients (direct recycling) or as detritus releasing mineral nutrients during decomposition (indirect recycling) (Vanni, 2002; Zou et al., 2016). Direct recycling is faster than indirect recycling because decomposition is required before the return of nutrients to the mineral pool, leading to increased primary production (Zou et al., 2016). Increasing the fraction of direct recycling should amplify the enrichment effect by accelerating the recycling. Increasing the decomposition rate of detritus should have a similar effect, especially if direct recycling does not prevail.

To study the consequences of nutrient cycling on food web response to nutrient enrichment and explore the mechanisms involved, we extended the recent food web modelling approach based on allometric relations with species body mass (*e.g.* Brose et al. (2006b); Heckmann et al. (2012); Schneider et al. (2016); Wang and Brose (2017)) by integrating basic aspects of nutrient cycling in this framework. Species body mass is linked to fundamental species traits such as metabolic or growth rates (Yodzis and Innes, 1992; McCann et al., 1998; Brown et al., 2004b) and it is also a good predictor of trophic interactions in ecosystems (Williams and Martinez, 2000; Petchey et al., 2008). Models parametrised with such allometric relations have been increasingly used to study food web dynamics and stability, especially because they tend to reproduce well, even though still simplified, observed patterns and dynamics of complex food webs (Boit et al., 2012; Hudson and Reuman, 2013). This framework thus offers a good opportunity to include nutrient cycling to food web models. To disentangle the mechanisms by which nutrient cycling affects food web stability (defined by species persistence and time variability of biomass dynamics), we assessed and compared the respective impact of nutrient cycling through the addition of mineral resources and the addition of feedback loops in both a complex food web and a food chain. These aspects were critical to answer the following questions: How nutrient cycling affect the overall nutrient availability in ecosystems and thus interact with the paradox of enrichment? Can the addition of feedback loops by nutrient cycling change the effects of the paradox of enrichment on species dynamics? Do the relative importance of direct and indirect nutrient cycling and the decomposition rate modulate these effects?

## Material and methods

### General description of the model

We developed a food web model including basic aspects of nutrient cycling by combining food web, allometry and stoichiometric theories (Fig. 1). Following classical allometric food web models (Brose, 2008; Heckmann et al., 2012), that are based on carbon flows, species biological parameters and trophic interactions scale with species body mass. Our model adds two major abiotic compartments, mineral nutrients (*e.g.* mineral nitrogen pool) and detritus (dead organic matter), to food web dynamics. Since detritus and mineral nutrient compartments are expressed in mass of nutrient whereas species compartments are expressed in mass of carbon, stoichiometry rules ensure the conversion between carbon flows and nutrient flows between the biotic and abiotic compartments and account for species stoichiometric homoeostasis in the food web. Nutrients are either directly recycled (species excretion of mineral nutrients directly available for primary producers) or indirectly recycled (species excretion of detritus releasing mineral nutrients through decomposition). All stocks are expressed for an arbitrary unit of habitat either a surface or a volume. The model is parametrised for nitrogen, but could be applied to other limiting nutrients such as phosphorus.

**Figure 1.**
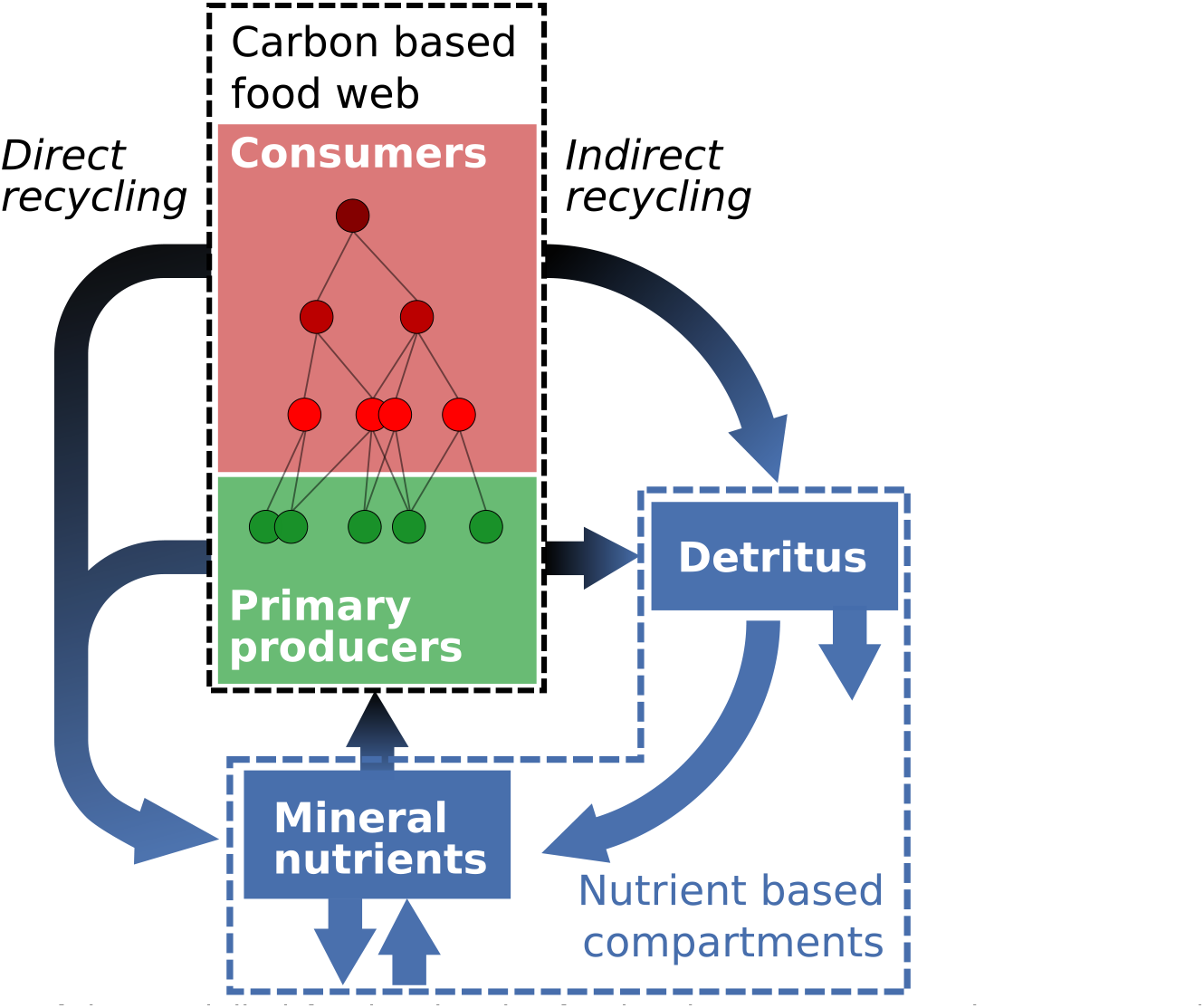
Schematic diagram of the modelled food web. The food web contains several primary producers and consumers forming a complex interaction network. It also includes two non-living compartments: mineral nutrients and detritus. Each organism excretes nutrients either directly as mineral nutrients (arrows on the left), or indirectly through the production of degradable detritus (arrows on the right). Stoichiometric rules ensure the conversions between the carbon based food web and the nutrient based compartments.

### Predator-prey interactions in the allometric food web model

For modelling food web dynamics, one needs to model both the structure of the food web (*i.e.* who eats whom) and the population dynamics within the food web. To define trophic interactions between species (*i.e.* food web structure), we took inspiration from the approach of the allometric diet breath model (ADBM, Petchey et al. (2008); Thierry et al. (2011)) because it predicts well trophic interactions in real food webs from species body mass and does not require additional assumptions on food web connectance (Petchey et al., 2008). To each of the 50 initial species is attributed a value *c* drawn uniformly in the interval [*−*5; 1]. Then, their body mass *M* is calculated as follow:

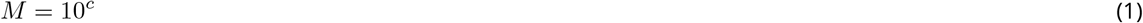

The five smallest species are defined as primary producers, the other as consumers. The diet of consumers depends on the profitability of each prey based on prey handling (*i.e.* the lower is the handling time, the more profitable is the prey). We derive the expression of the mass specific handling time *h*_*ij*_ of species *j* by the consumer *i* from Petchey et al. (2008) and Thierry et al. (2011) and (see Appendix S1 in the supporting information):

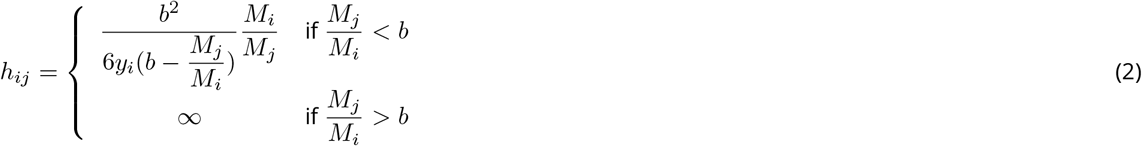

With *y*_*i*_ the maximum ingestion rate (see equation (4e)), *M*_*j*_ the body mass of the prey, *M*_*i*_ the body mass of the consumer and *b* the maximum prey-predator body mass ratio above which the prey cannot be eaten. The handling time function against prey body mass is U-shaped (see Fig. S1-1 in the supporting information), handling time being minimal when prey body mass is equal to *b*/2 × *M*_*j*_. We consider that predators can only interact with preys within the body-mass interval [0.1*bM*_*i*_, *bM*_*i*_] with *b* < 1 (*i.e.* predators are always larger than their prey) as the handling time increases exponentially out of this interval. Thus, the structure of the food web (*e.g.* number of trophic levels, see Fig. S2-2C and D in supporting information) is shaped by species biomass distribution and body mass ratios between species.

The predator-prey dynamics follow previous allometric food web models (Brose, 2008; Heckmann et al., 2012). The respective equations for primary producers (equation 3a) and consumers (equation 3b) are:

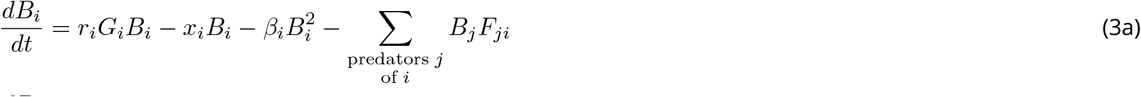

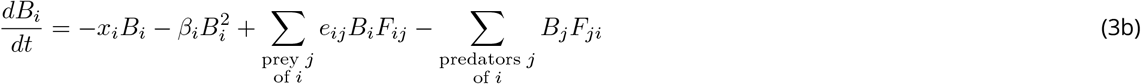

In these equations, *B*_*i*_ is the biomass of population *i*, *G*_*i*_ is the nutrient-dependent growth rate of primary producers, *r*_*i*_ is the mass-specific maximum growth rate of primary producers, *x*_*i*_ is the mass-specific metabolic rate, *β*_*i*_ is the density dependent mortality rate (ensuring a reasonable species persistence, see Fig. S3-4 and S3-5A in the supporting information) and *e*_*ij*_ the assimilation efficiency of species *j* by species *i*. Primary producer growth rates *r*_*i*_, species metabolic rates *x*_*i*_, density dependent mortality rates *β*_*i*_, consumer attack rate *a*_*i*_ and maximum ingestion rates *y*_*i*_ (involved in handling time parametrisation, see equation (2)) are defined as functions of species body masses, according to the allometric quarter-power laws as described by Yodzis and Innes (1992) and Brown et al. (2004b):

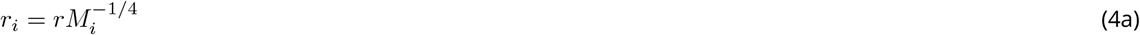

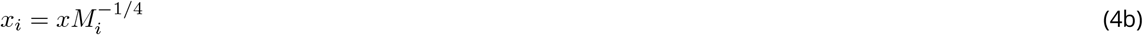

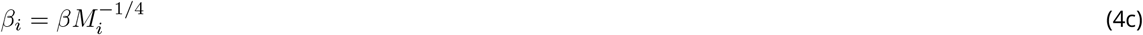

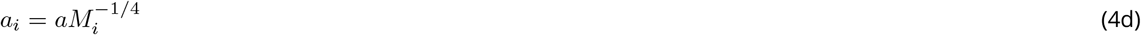

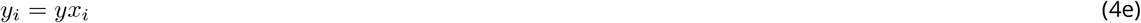

With *M*_*i*_ the body mass of species *i* and *r*, *x*, *β*, *a* and *y* being allometric constants (Table 1, see also Appendix S1 in supporting information).

**Table 1.** Table of variables and parameters (below the horizontal separation). *v* represents a generic metric of space (*e.g.* that could represent litres or square meters). Indeed all the parameters depending on space are set arbitrarily and thus we do not need to specify a particular unit of space. The values of *β* and *a* are set arbitrary to ensure a reasonable species persistence and and time variability of species biomasses (See Fig. S3-5 in the supporting information). *K* and *ℓ* are set to ensure a maximal persistence at *I* ∼ 50.

|  | Value and units | Description | Reference |
| --- | --- | --- | --- |
| $B_i$ | $kg.v^{-1}$ | Species biomass (carbon) of species $i$ | Variable (equation 3a, 3b) |
| $N$ | $kg.v^{-1}$ | Mineral nutrient stock (nitrogen) | Variable (equation 7a) |
| $D$ | $kg.v^{-1}$ | Detritus stock (nitrogen) | Variable (equation 7b) |
| $\omega_{ij}$ | Dimensionless | Preference of predator $j$ for prey $i$ | Variable (equation 6) |
| $r$ | $0.87\ kg^{1/4}.year^{-1}$ | Growth rate allometric constant | Binzer et al. (2012) |
| $x$ | $0.12\ kg^{1/4}.year^{-1}$ | (primary prod.) Metabolic rate | Brose (2008) |
| | $0.27\ kg^{1/4}.year^{-1}$ | (consumer) allometric constant | |
| $y$ | 8 | Maximum ingestion rate factor | Brose (2008) |
| $\beta$ | $0.001\ v.kg^{-3/4}.year^{-1}$ | Density dependent mortality rate allometric constant | Arbitrary (Fig. S3-4 and S3-5) |
| $a$ | $0.1\ v.kg^{-3/4}.year^{-1}$ | Attack rate allometric constant | Arbitrary (Fig. S3-5) |
| $h_{ij}$ | year | Handling time | Equation (2), Appendix S1 |
| $e_{ij}$ | 0.45 (herbivore) | Assimilation efficiency of species $j$ | Yodzis and Innes (1992) |
| | 0.85 (carnivore) | eaten by species $i$ | |
| $q$ | 1 | Hill exponent | Brose et al. (2006b) |
| $A$ | 0.01 | Adaptive rate | Arbitrary (Fig. S3-6) |
| $b$ | 0.05 | Max prey-predator body mass ratio | Brose et al. (2006a) |
| $\alpha_i$ | 6.6 (primary prod.) | Constant carbon to nutrient ratio | Anderson (1992) |
|  | 5 (consumer) |  |  |
| $K_i$ | $10\ kg.v^{-1}$ | Half saturation of nitrogen uptake | Arbitrary (Fig. S3-3) |
| $\ell$ | $0.2\ year^{-1}$ | Leaching rate | Arbitrary (Fig. S3-3) |
| $M_i$ | kg | Body mass of species $i$ | Log uniform in $[10^{-5}, 10]$ |
| $I$ | $kg.v^{-1}.year^{-1}$ | External mineral nutrient input | $[0, 400]$ |
| $d$ | Dimensionless | Decomposition rate of detritus | $[0, 1]$ |
| $\delta$ | Dimensionless | Fraction of direct recycling | $[0, 1]$ |

*F*_*ij*_ represents the contribution of species *j* in the eaten biomass per unit species *i* biomass and follows a Holling functional response:

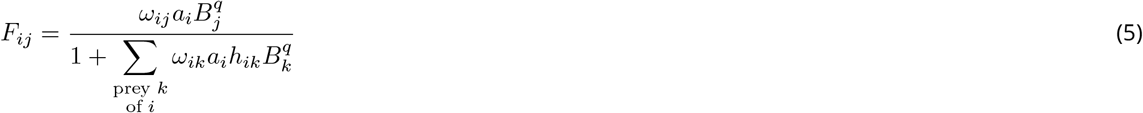

*B*_*j*_ represents the biomass of the prey *j*, *q* is the Hill exponent (the functional response is of type II if *q* = 1 or type III if *q* > 1), *a*_*i*_ is the attack rate of consumer *i* and *h*_*ik*_ is the handling time of *k* by consumer *i*. *ω*_*ij*_ is the preference of *i* for the prey *j*. We chose here to model preferences as time variables and not as fixed parameters according to the adaptive foraging theoretical framework (results with preferences as fixed parameters are available in Fig. S3-6 the in supporting information). Adaptive foraging is indeed an important aspect of predator-prey interactions (*e.g.* predator foraging efforts depend on prey availability) and it strongly affects food web dynamics (Kondoh, 2003; Uchida and Drossel, 2007; Heckmann et al., 2012). The dynamics of foraging efforts were modelled through changes over time of the consumer preferences *ω_ij_* according to the following equation:

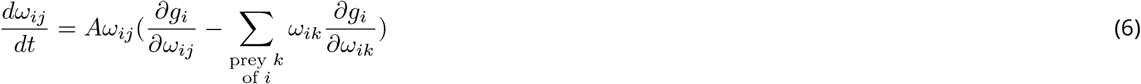

Here, *A* represents the adaptive rate of the diet preference and *g*_*i*_ the total growth rate of species *i* defined such as 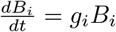. The initial value of *ω*_*ij*_ is set assuming a uniform distribution among preys and during the simulation, the *ω*_*ij*_ are rescaled after the resolution of equation 6 to keep the relation 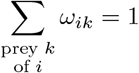 true at each time step.

### From a carbon-based food web model to an ecosystem model including nutrient cycling

To expand the classical food web model to take fundamental aspect of nutrient cycling into account, we model the dynamics of two abiotic compartments, mineral nutrients *N* and detritus *D*. These compartments are described as masses of nutrient while species biomasses are based on carbon in the food web model. We use species carbon to nutrient ratios (C:N) *α*_*i*_ to convert carbon flows into nutrient flows (and vice versa). For simplicity, we assume the *α*_*i*_ to be constant over time. Please note that we could have expressed directly the species biomasses in nutrient instead (as in Zou et al. (2016)), without changing the model behaviour. However, we chose to keep species biomasses based on carbon to relate more clearly our equations with classical allometric food web models. The dynamics of nutrients in the mineral and detritus compartments are described by:

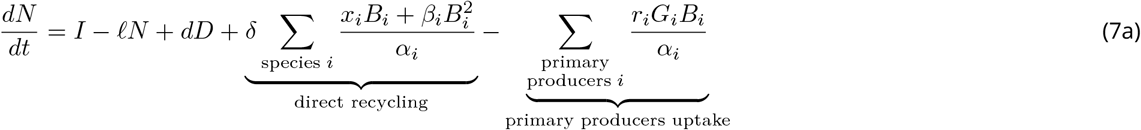

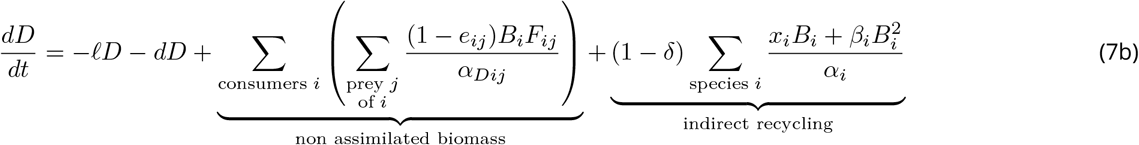

We consider an open ecosystem where *I* is the constant external input of nutrients (*e.g.* through erosion or atmospheric deposition) and *ℓ* is the rate of loss of mineral nutrients and detritus (*e.g.* through leaching, sedimentation). The nutrient dependent growth rate of primary producers is expressed as (DeAngelis, 1980; DeAngelis et al., 1989):

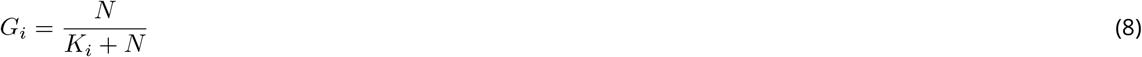

*K*_*i*_ is the half saturation constant of nutrient uptake of primary producer *i*. The nutrient uptake by primary producers (expressed as a nutrient flow) is calculated by dividing the growth rate of primary producers (expressed as a carbon flow) by their C:N ratio *α*_*i*_. Detritus are decomposed at a constant rate *d*. Organisms release nutrients through excretion and mortality to the detritus and mineral nutrient pools (Fig. 2B). A fraction *δ* of these nutrients is released in their mineral form (urine for instance) while the remaining fraction is released as dead organic matter (detritus like feces, dead bodies, litter fall...) as in Zou et al. (2016). We assume that the nutrients contained in the non-assimilated biomass (1 − *e*_*ij*_) go in the detritus compartment. The amount of nutrients released by species in the food web depends on their C:N ratio *α*_*i*_. The carbon to nutrient ratio of non-assimilated biomass *α*_*Dij*_ depends on both the C:N ratio of the prey *j* and of the consumer *i* (calculation detailed in Appendix S1 of supporting information):

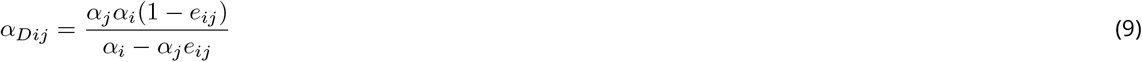

**Figure 2.**
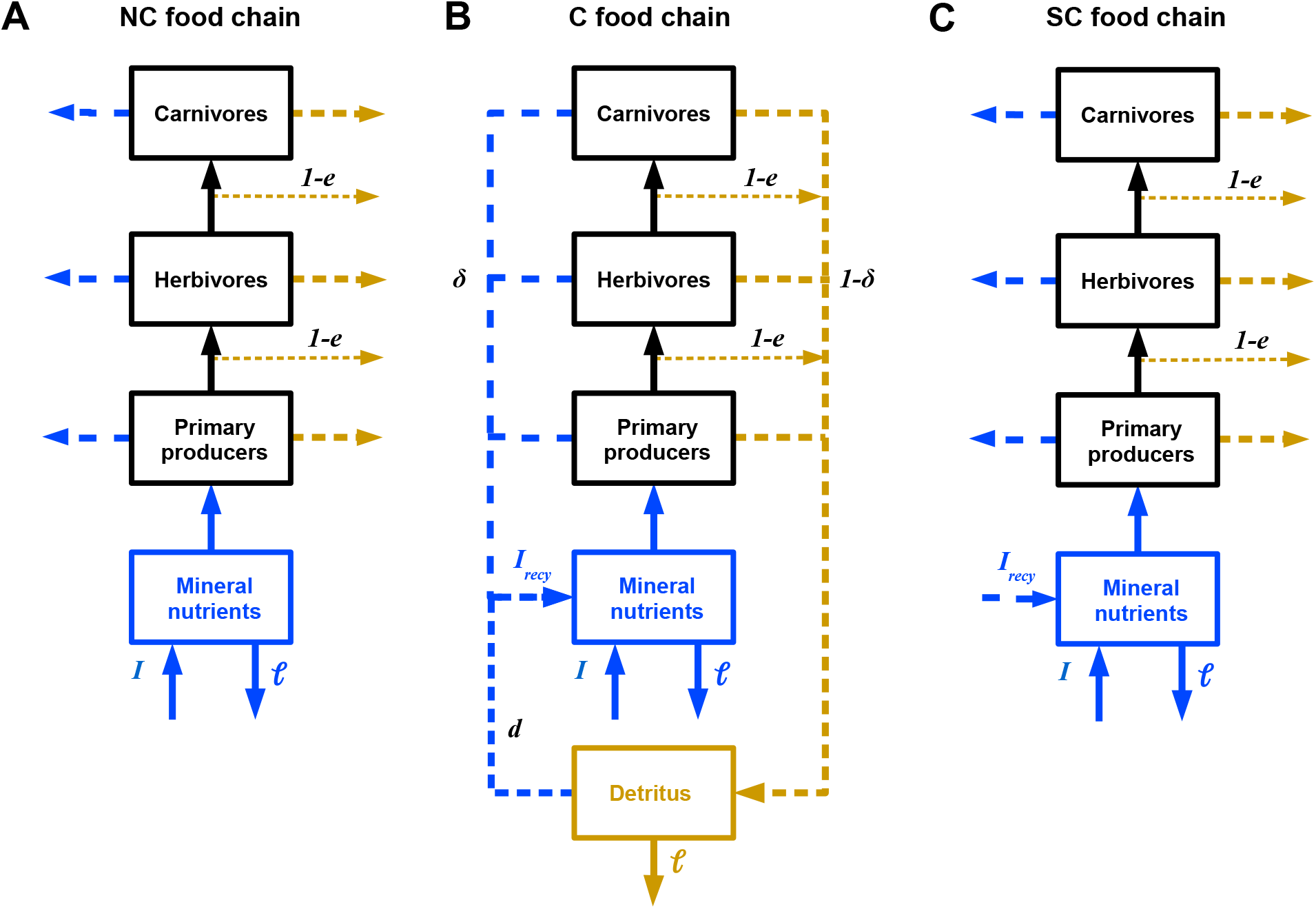
Diagram of the general structure of our models with and without nutrient recycling feedback loops. Food chains are represented for more simplicity but these three models are valid for food webs as well. The dotted arrows represent nutrient cycling (nutrient flux in blue, detritus in brown). **A)** NC model. Food chain without nutrient cycling. **B)** C model. Food chain with nutrient cycling. A fraction *δ* of nutrients is excreted as mineral nutrients (direct recycling on the left) and a fraction 1 − *δ* plus a fraction 1 − *e* of non ingested biomass are excreted as detritus (indirect recycling on the right). The total nutrient input *I*_*tot*_ in the pool of mineral nutrients is the sum of the external nutrient input *I* and the recycled nutrient *I*_*recy*_. **C)** SC model. Food chain without nutrient cycling but with a nutrient input corrected by *I*_*recy*_. The resulting food chain does not have the feedback loop induced by nutrient recycling but has an equivalent nutrient availability as in the food web with nutrient cycling. Note that the first version of our model (NC) is based on the C model where *I*_*recy*_ is set to 0.

### Assessing nutrient cycling effects on stability

Stability was assessed by two measures: species persistence and the average coefficient of variation of species biomass (CV) weighted by the relative biomass of each species. To investigate the effects of nutrient cycling on food web dynamics and disentangle effects due to enrichment from effects due to presence of additional loops, each food web was studied for three configurations of nutrient cycling (Fig. 2). (1) No nutrient cycling with the fraction of direct recycling *δ* and the decomposition rate *d* set to zero. This corresponded to the dynamics obtained with classic allometric food web models and will be referred as the NC model (No Cycling) (Fig. 2A). (2) With nutrient cycling with the fraction of direct recycling *δ* and the decomposition rate *d* strictly positive (Fig. 2B). This food web was referred as the C model (Cycling). (3) No nutrient cycling but the enrichment effect of nutrient cycling was simulated (Fig. 2C). This food web was referred as the SC model (Simulated Cycling). In this last case, we removed the potential effect of the temporal coupling between higher trophic levels and the basal resource due to the presence of recycling loops while keeping the additional inputs of nutrients associated with nutrient cycling. To simulate the enrichment effect of nutrient cycling, we replaced the basal nutrient input by the total nutrient input *I*_*tot*_:

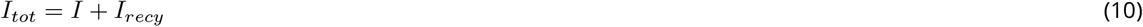

Where *I* is the external nutrient input and *I*_*recy*_ is the average quantity of recycled nutrients (directly and indirectly) calculated over the last 1000 years of the simulation in the C model. Thus, in the SC model, *I*_*recy*_ was constant over time because recycling (both direct and indirect recycling) was not explicitly modelled. In the C model, these nutrient subsidies varied over time as they directly originated from the direct and indirect recycling loops and thus depended on temporal variations of species biomasses and detritus in the ecosystem.

### Simulations

All the parameters, their units and their values as used in the simulations are given in table 1. The sensitivity of the results to arbitrarily set parameters is given in Appendix S3 in supporting information. The simulations were performed with *C* + + and the *GSL* ode solver using the Runge-Kutta-Fehlberg (4, 5) method with an adaptive time step and an absolute accuracy of 10^*−*6^. Simulations were run as follow: first, 50 species are attributed a body mass (the five smallest being primary producers) and trophic links were set depending on predator-prey body mass ratios (see equation 2). We did not seek for food webs with our 50 species linked by trophic interaction, thus consumer without prey got extinct during simulations. Then, simulations were run a for 9000 years to let the ecosystem reach a steady state. We kept in our results all resulting food webs even when some of the initial 50 species got extinct (see Fig. S3-1C in the supporting information). Species were considered as extinct if their biomass fell below 10^*−*30^ kg.v^*−*1^ and consumers without prey got extinct. After this preliminary phase, outputs were recorded for 1000 years. Species persistence was measured as the ratio of the final number of species at *t* = 10, 000 to the initial number of species at *t* = 0. The CV was the ratio of the standard deviation to the mean of species biomass or recycled quantity of nutrients over time, calculated for the 1000 last years of each simulation. Each combination of parameters was tested for 100 different food webs (*i.e.* different randomly drawn sets of species body mass), each of these food webs being simulated in the three configurations of nutrient cycling (*i.e.* for the NC, C and SC models). To implement the SC model, we recorded the density of each compartment in the simulation of the C model at *t* = 9000 as well as the averaged quantity of recycled nutrient *I*_*recy*_ for the last 1000 years. We then ran corresponding food web simulations for the SC model (*i.e.* with *δ* = *d* = 0 and *D* = 0) for 1000 years with initial densities and a nutrient input *I* respectively set equal to the densities and *I*_*tot*_ recorded in the C model.

In each simulation for complex food webs, there were initially 50 species and their initial biomasses were set at 10 kg.v^*−*1^ for primary producers and at 5 kg.v^*−*1^ for consumers (v is an arbitrary metric of space, see table 1). Initial quantities of nutrients in the mineral nutrients and detritus pools were set at 10 kg.v^*−*1^.

## Results

### Overall effects of nutrient cycling on food web dynamics

Nutrient cycling contributes to an important part of the total mineral inputs of nutrients in the food web, and its contribution increases with the levels of external inputs of nutrients (Fig. 3A), in parallel with variations of primary and secondary productions (see Fig. S2-4B in supporting information). In this study case, nutrient cycling always represents larger inputs of nutrients to the food web than external inputs (Fig. 3A). At low nutrient enrichment levels, consumers are responsible for a significant part of direct recycling (see Fig. S2-3A in supporting information) and indirect recycling as they release more than 50% of detritus (see Fig. S2-3B in supporting information). However, at high nutrient enrichment levels, the quantity of nutrient directly recycled by consumers stops increasing while the total quantity of nutrient recycled still increases linearly with the external nutrient input *I* due to a large increase in the quantity of nutrient directly cycled by primary producers. Similarly, consumer biomass production is relatively important at low external nutrient input *I* while primary production is dominant and increases linearly with *I* at high inputs (see Fig. S2-4B in supporting information).

**Figure 3.**
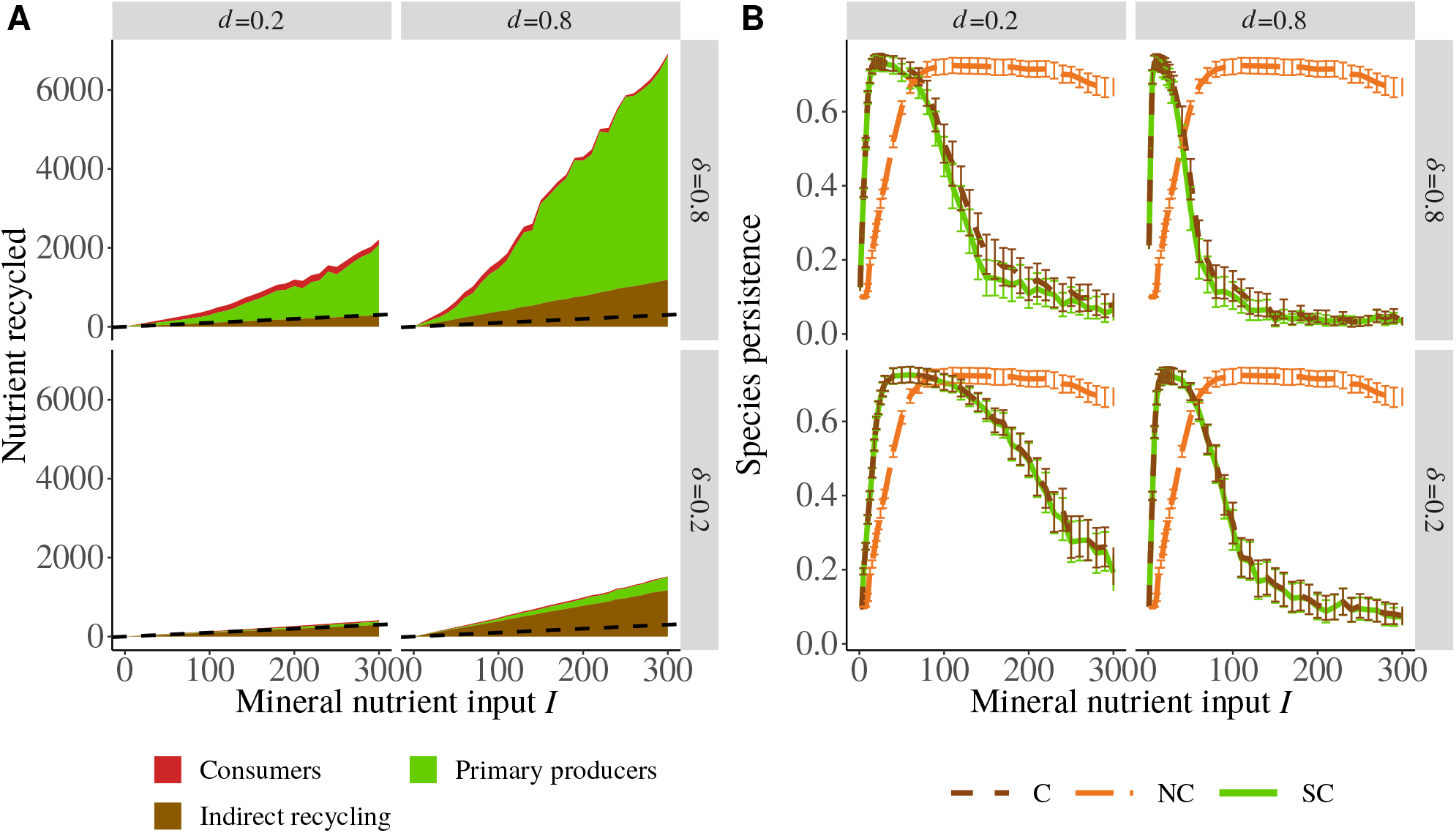
Food web responses to a nutrient enrichment gradient (increasing *I*) as a function of recycling parameters *d* (decomposition rate) and *δ* (proportion of direct recycling). **A)** Average quantity of nutrients recycled directly by consumers (red), primary producers (green) and indirectly recycled (brown) in C foo webs. The dashed line (with a slope equal to 1) represents cases where the average quantity of recycled nutrients is equivalent to the external nutrient input *I*. Only food webs where at least one species persists are kept. **B)** Effects of nutrient cycling on species persistence (proportion of surviving species at the end of simulations). The brown dashed curve represents the C food webs with nutrient cycling (*δ* > 0, *d* > 0), the orange long-dashed curve represents the NC food webs without nutrient cycling and the green solid curve represents the SC food webs without nutrient cycling but with a mineral nutrient input simulating the enrichment effect of nutrient cycling in the C food web. 100 different food webs randomly generated are tested for each combination of parameters. Outputs are then averaged and the error bars represent the confidence intervals of the mean.

Nutrient cycling affects the food web response to nutrient enrichment (*i.e.* external nutrient inputs *I*). First, it affects the relationship between species persistence and nutrient enrichment (Fig. 3B). In food webs with and without nutrient cycling, persistence follows a hump-shaped relationship with external nutrient input *I*: first there is a sharp increase of the persistence for low nutrient inputs, then a plateau with maximum persistence and finally a decrease of the persistence for high nutrient inputs. However, maximum persistence is reached for lower input values and effects of enrichment are sharper for the case with nutrient cycling (C) than for the case without nutrient cycling (NC). These sharp changes in species persistence along the gradient of nutrient enrichment are paralleled by strong changes in food web maximum trophic level with an increase and then a decrease of the maximum trophic level with increasing external nutrient input *I* (see Fig. S2-2B and S2-2C in supporting information).

Second, nutrient cycling affects the relation between the average coefficient of variation (CV) of the species biomass and nutrient enrichment (Fig. 4A). As for species persistence, the average species biomass CV first increases and then decreases with increasing external mineral nutrient inputs. This decrease is due to the existence of food webs where only primary producers survived, leading to constant biomasses (see Fig. S2-2D in supporting information). As for species persistence (Fig. 3B), maximum biomass CV is reached for lower input values without nutrient cycling (NC) than for the case with nutrient cycling (C). The CV of the quantity of recycled nutrients (Fig. 4B) and of the nutrient stock (Fig. S2-2A in supporting information) follow a hump-shaped relation with external nutrient input *I* but the temporal variability of the quantity of recycled nutrients is about 25 times smaller than the CV of species biomass (see also Fig. S2-1A and S2-1B in supporting information).

**Figure 4.**
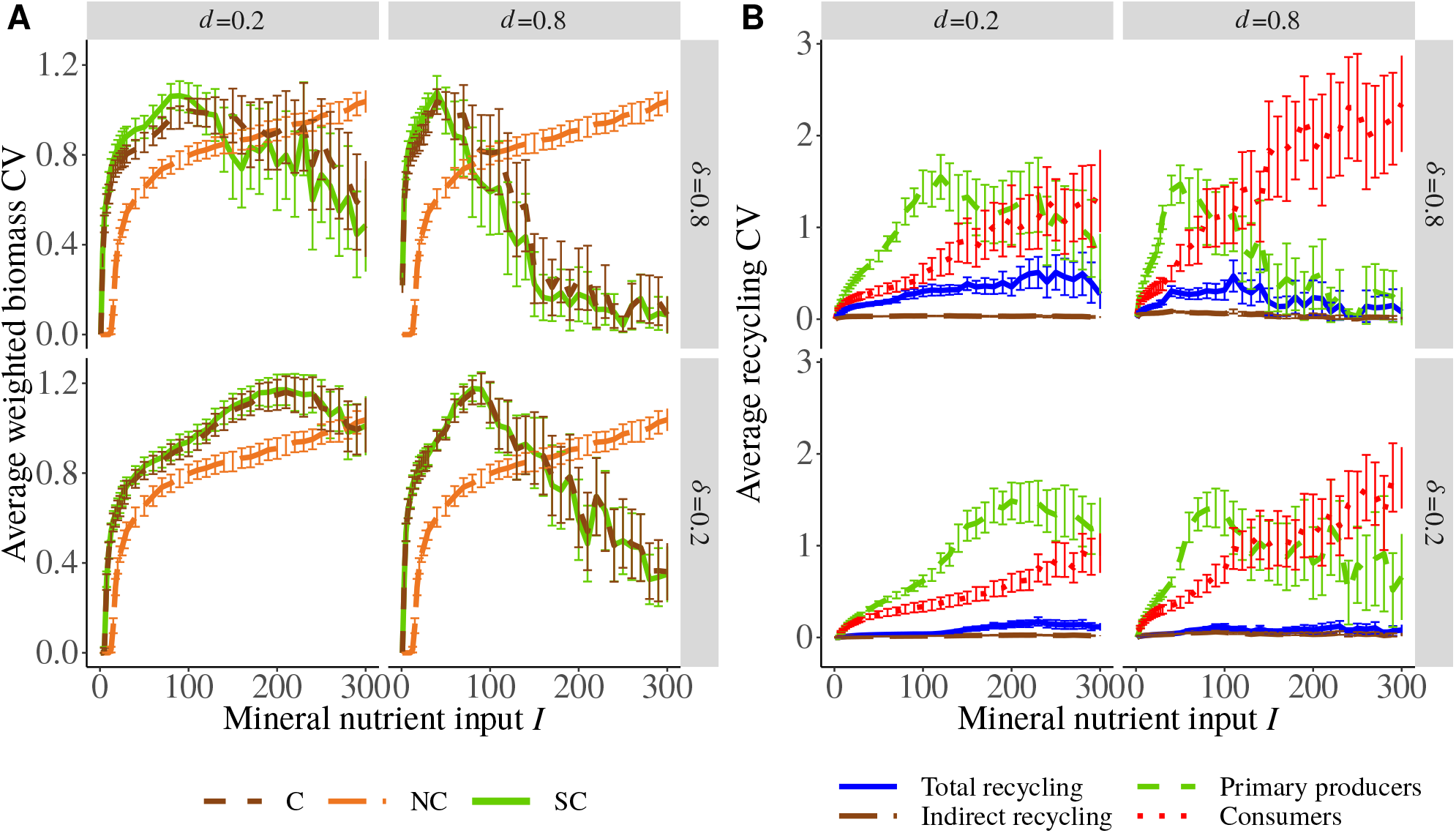
Responses of species and recycling temporal variability to a nutrient enrichment gradient (increasing *I*) as a function of recycling parameters *d* (decomposition rate) and *δ* (proportion of direct recycling). **A)** Effects of nutrient cycling on the average weighted coefficient of variation (CV) of species biomass. CVs are weighted by the contribution of the biomass of each species to the total biomass (*CV*_*i*_ × *B*_*i*_/*B*_*tot*_). **B)** Average CV of nutrient recycled in total (solid blue), indirectly recycled (long-dashed brown), directly by primary producers (dashed green) and directly by consumers (dotted red) in C food webs. Only food webs with at least on persisting consumer (and consequently at least one persisting primary producer) are kept for calculations. 100 different food webs randomly generated are tested for each combination of parameters and the error bars represent the confidence intervals of the mean.

### Overall influence of the recycling parameters

Increasing the decomposition rate *d* and the fraction of directly recycled nutrients *δ* always increases the quantity of recycled nutrients in the food web (Fig. 3A), leading to greater inputs of nutrients through recycling than external inputs. *d* and *δ* both increase the quantity of directly recycled nutrients while only *d* increases the quantity of indirectly recycled nutrients. In fact, the detritus stock does not depend on recycling parameters (see Fig. S2-4D in the supporting information) and the mineral nutrient stock is always controlled by primary producers and their quantity is negligible compared to detritus. Thus, the external nutrient input *I* is mainly balanced by the loss from detritus *ℓD*, leading at equilibrium to *D*^∗^ = *I/ℓ* that does not depend on *d* and *δ*. Therefore, the average quantity of indirectly recycled nutrients is equal to *dD*^∗^.

Both *d* and *δ* affect the relationship between external nutrient input *I* and species persistence or species biomass CV (Fig. 3B and 4A). At high *d* and *δ*, the increase and decrease of species persistence and biomass CV with increasing nutrient input *I* are sharper. However, the general response of the food web remains qualitatively unchanged. In addition, unlike *d*, high values of *δ* amplify the destabilising and stabilising effects of feedback loops on primary producer (Fig. 5A) and consumer (Fig. 5B) dynamics respectively (this aspect is detailed in the following). Finally, increasing *δ* increases the CV of the total quantity of recycled nutrients (Fig. 4B) by increasing the contribution of direct recycling (Fig. 3A) that has a higher CV than indirect recycling.

**Figure 5.**
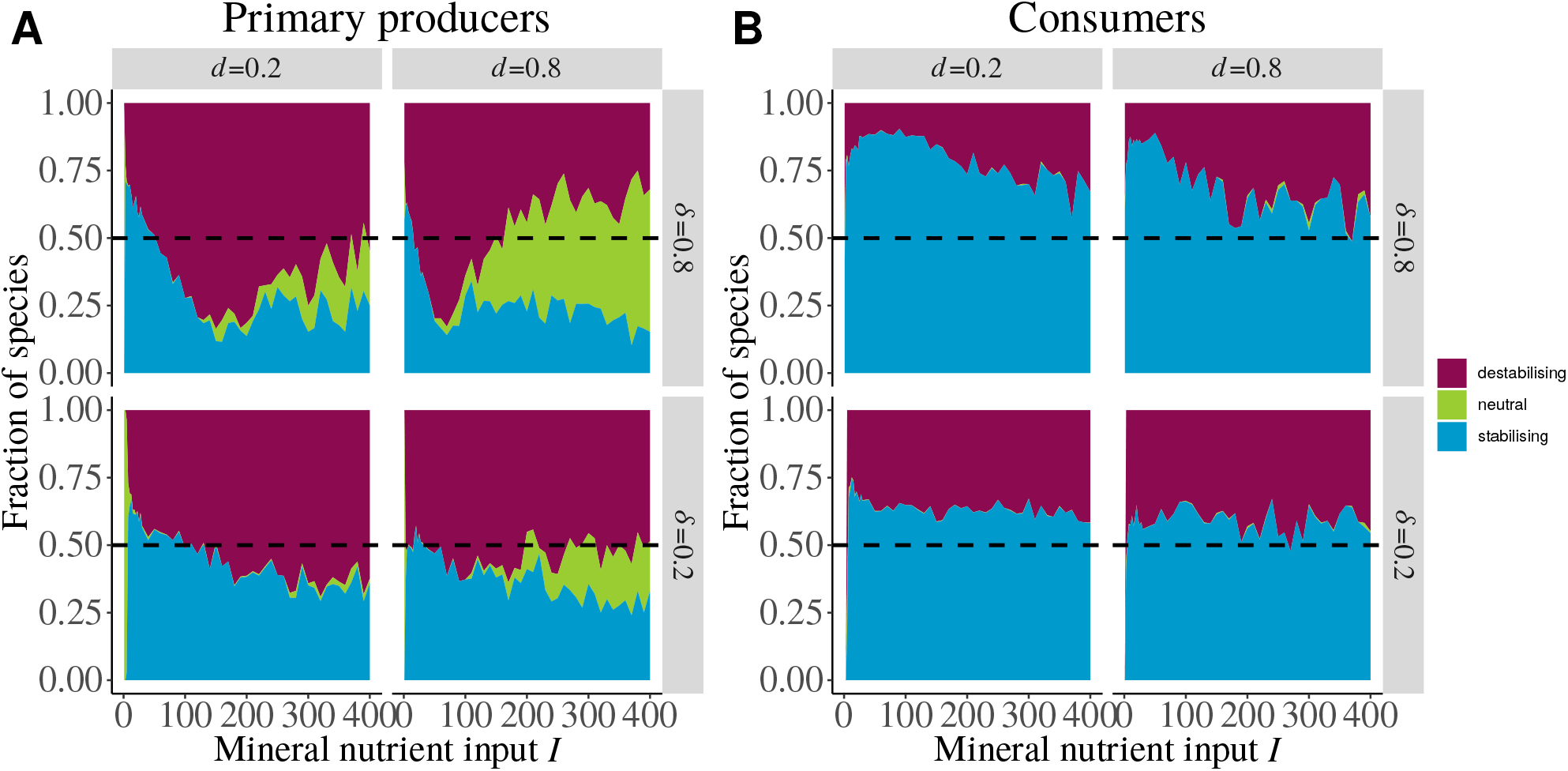
Effect of nutrient cycling on biomass CV at species level. For each combination of parameters, the biomass CV of persisting species is compared between C ans SC food webs with the same species. If the CV is higher in the SC food web (without nutrient cycling but with a mineral nutrient input simulating the enrichment effect of nutrient cycling) than in the C food web (with nutrient cycling) with a threshold set at 10^*−*4^, then nutrient cycling feedback loops have a stabilising effect on dynamics. We have the same conclusion if the species get extinct in the SC food web and not in the C food web. The fraction of stabilised or destabilised **A)** primary producers and **B)** consumers among all simulated food webs gives the overall effect of nutrient cycling feedback loops at species level.

### Effects of nutrient cycling: enrichment vs feedback loop

The comparison between the case with nutrient cycling (case C) and the case without nutrient cycling but with a nutrient input simulating the enrichment effect of nutrient cycling (case SC) allows to separate the effects of nutrient cycling due to enrichment from those due to the creation of additional feedback loops from each trophic levels to the bottom of the food web. When we model food web dynamics without nutrient cycling but including the enrichment effect of nutrient cycling (*i.e.* SC case), the overall relationships between external nutrient inputs and species persistence or biomass CV are similar to those observed in presence of nutrient cycling (Fig. 3B and 4A). Indeed, the curves corresponding to C and SC strongly overlap. Most of the effects of nutrient cycling are thus due to an enrichment effect caused by recycled nutrients. However, curves do not overlap perfectly for *δ* = 0.8, the average species persistence is higher and species biomass CV is lower (at low nutrient input *I*) in food webs with nutrient cycling (C model). This is emphasised in Fig. 5B (see also Fig. S2-5B-D in supporting information) where more than 50% of consumers from all simulated food webs put together have a lower CV in C food webs than in SC food webs (for *δ* = 0.8 and a low *I*) while primary producers tend to have a higher biomass CV (Fig. 5A). For *δ* = 0.2, less primary producers and consumers have respectively higher and lower biomass CVs in SC food webs but the trend seen for *δ* = 0.8 is still visible. The decomposition rate *d* weekly changes the fractions of species stabilised or destabilised by feedback loops compared to *δ*. It only increases the fraction of primary producers that are not affected (Fig. 5A), probably because of the extinction of higher trophic levels (see also Fig. S2-2B in supporting information) for high external nutrient inputs. Indeed, this leads to food webs where only primary producers persist with constant biomasses whatever the presence or not of feedback loops.

## Discussion

By integrating nutrient cycling and trophic dynamics, our food web model allows to better link population dynamics and ecosystem functioning. Our food web model highlights that nutrient cycling strongly interacts with the paradox of enrichment following two main mechanisms. First, nutrient cycling effects are mostly due to the consecutive increased nutrient availability that promotes species persistence at low nutrient inputs but leads to species extinctions (characteristic of the paradox of enrichment) at high level of nutrient inputs. Second, feedback loops from each species to the bottom resource generated by nutrient cycling only weakly affect species biomass temporal variability beyond effects associated with nutrient enrichment. Recycling loops tend to slightly dampen the destabilising effect of nutrient enrichment due to nutrient cycling on consumer dynamics while they have the opposite effect on primary producer dynamics. These results are thoroughly discussed below and their sensitivity to the parameters (Table 1) is tested in Appendix S3.

### Paradox of enrichment in our food web model

In agreement with previous food web studies (Rall et al., 2008; Binzer et al., 2016), we observe two contrasting responses of species diversity and food web stability to increased external nutrient inputs. While higher nutrient availability consistently increases the temporal variability of species biomasses, it also increases species persistence in nutrient poor ecosystems (*i.e.* low external nutrient inputs) but decreases persistence at high inputs of nutrients. The increase in persistence at low nutrient inputs is likely due to the increased persistence of species at higher trophic levels (Fig. S2-2B and S2-2C in the supporting information). Higher trophic levels are known to require a sufficient ecosystem productivity (limited by nutrient availability) to meet their energetic requirement and persist (*e.g.* Oksanen et al. (1981); Abrams (1993); Leibold (1996)), which can explain why increased persistence is only found in our case for nutrient poor ecosystems. The observed increase in the amplitude of species biomass oscillations (*i.e.* increase of species CVs that destabilises species dynamics, see Fig. 4A) with increasing nutrient inputs is typical of the well-known paradox of enrichment (Rosenzweig, 1971; DeAngelis, 1992; Roy and Chattopadhyay, 2007; Rip and McCann, 2011). Such destabilising effects of nutrient availability on species dynamics might explain the decrease in species persistence we observe at high levels of nutrient inputs. Large oscillations of species biomass (Fig. S2-1A in the supporting information) caused by nutrient enrichment likely trigger species extinctions as their biomass might reach the extinction threshold value (Fig. S3-2 in the supporting information). This counteracts the positive effect of nutrient enrichment on persistence at low nutrient levels and results in an hump-shaped relationship between species persistence and nutrient enrichment. Thus, parameters determining the occurrence of limit-cycles in complex food webs should strongly determine food web response to increased external nutrient inputs as well as nutrient cycling. For instance, the scaling of the attack rate with predator and prey body masses strongly determines the occurrence of limit cycles (Pawar et al., 2019) and varies a lot between studies (Rall et al., 2008; Pawar et al., 2012). However, such differences do not change our main results as the C and SC models respond similarly whatever the values of our scaling constants (Fig. S3-5 in the supporting information). In accordance with our model results, the paradox of enrichment has been found in complex food web models with type II functional responses (Rall et al., 2008; Binzer et al., 2016). In case of type III functional responses (Fig. S3-7 in the supporting information) or high intraspecific density dependence regulation (Fig. S3-4 in the supporting information), where no such destabilising effects occur (see also Rall et al. (2008)), our results show that persistence does not decline at high levels of nutrient availability. Adaptive foraging, as included in our study, does not affect the occurrence of the paradox of enrichment (Fig. S3-6 in the supporting information) as already observed by Mougi and Nishimura (2008) in a one predator-two prey model. Our results remain indeed qualitatively the same with and without adaptive foraging.

### Nutrient cycling and enrichment effects

Our results show that nutrient cycling mainly interacts with the paradox of enrichment through its impacts on nutrient availability in ecosystems. Indeed, the quantity of recycled nutrients is, in our model, from one to more than ten times larger than the external input of mineral nutrients (depending on the decomposition rate *d* and the fraction of direct recycling *δ*). This ratio is consistent with the flows measured for the biogeochemical nitrogen cycle in natural ecosystems (Gruber and Galloway, 2008; Fowler et al., 2013). Thus, nutrient cycling strongly amplifies food web response to external nutrient inputs: the effects described in the previous section are qualitatively similar with and without nutrient cycling but they occur for lower inputs when nutrient cycling is present. Two main mechanisms rule the enrichment effect of nutrient cycling.

First, factors increasing the recycling speed (*i.e.* higher decomposition rate *d* and fraction of direct recycling *δ*) lead to species persistence and CV values that are obtained for increased levels of nutrient inputs in food webs with a slower nutrient cycling. They thus amplify the enrichment effect of nutrient cycling and also interact with the paradox of enrichment. In fact, nutrient cycling has been shown to increase the total amount of mineral nutrient circulating in the ecosystem and primary production (DeAngelis, 1980; de Mazancourt et al., 1998; Barot et al., 2007; Loreau, 2010). In our model, *d* and *δ* mainly rule the flows between compartments but weakly control compartment size. Indeed, the detritus compartment size does not depend on decomposition rate *d* (see Fig. S2-4D in the supporting information) and the total biomass is mostly related to external nutrient inputs and species persistence (see Fig. S2-4A in the supporting information). Thus, as compartment size does not depend on *d* and *δ*, nutrients released by detritus decomposition (*d*) or direct recycling (*δ*) must be compensated to keep the matter balance in the ecosystem. Given the absence of nutrient loss by the ecosystem from the species compartment and the small size of the mineral nutrient compartment (due to the control by primary producers), recycled nutrients must reflow in the species compartments. This leads to an increase of biomass production (see Fig. S2-4B in the supporting information), compensated by an increased mortality due to density dependent self-regulation (see Fig. S2-4C in the supporting information) that leads to an increased quantity of recycled nutrients that fuels biomass production and so on. This suggests that the impact of nutrient cycling partly arises in our models from complex interactions between the speed of recycling and nutrient losses (see Fig. S3-3 in the supporting information). These interactions should be further disentangled through new simulations manipulating independently rates of mineral nutrient and detritus loss that are set equal in our model while higher losses for mineral nutrients than for detritus would be more realistic, at least in terrestrial ecosystems. In addition, density dependent mortality seems to have a strong quantitative impact on nutrient cycling in our model. Although it does not affect the qualitative response of species persistence and biomass CV to nutrient enrichment (see Fig. S3-4 in the supporting information), it drastically increases the quantity of nutrients flowing out of the species compartment and then through the entire ecosystem. Other mechanisms limiting species biomass such as predator interference in the Beddington-DeAngelis functional response (Beddington, 1975; DeAngelis et al., 1975) decrease the net growth rate by reducing the resource uptake rate instead of increasing the death rate. As a consequence, such a mechanism would lead to reduced nutrient flows in the ecosystem, thus changing nutrient cycling. Such effects of population dynamics modelling on ecosystem functioning must be explored in future studies.

Second, the amount of recycled nutrients depends on food web structure (relative importance of the trophic levels and the food chain length) and strongly depends on primary production, which increases linearly with nutrient inputs (Loreau, 2010). In fact, nutrient uptake by producers necessarily balances nutrients recycled from all trophic levels at equilibrium (see Fig. S2-4B and S2-4C in the supporting information). At low nutrient inputs, consumers are the main contributors to nutrient cycling, in agreement with experimental and empirical studies (Vanni, 2002; Schmitz et al., 2010) (see Fig. S2-3 in the supporting information). While nutrients recycled per unit of biomass due to species metabolism are lower for consumers because of their larger body mass, consumers also strongly contribute to recycling through nutrient losses associated to feeding inefficiency. This is particularly true for herbivores whose assimilation efficiency is low (*e*_*ij*_ = 0.45) so that they produce a lot of detritus by consuming primary producers (see Fig. S2-3B in the supporting information). This is also emphasised by previous ecosystem models (Leroux and Loreau, 2010; Krumins et al., 2015). However, at high nutrient input, food webs are dominated by primary producers (see Fig. S2-4A in the supporting information) that become the main contributors to nutrient cycling. In such case, primary producers release large amounts of detritus and nutrients due to high metabolic rates and large density dependent mortalities (see Fig. S2-4C in the supporting information).

Food web structure influences nutrient cycling through other already identified mechanisms pertaining to the quality of the produced detritus that are not included in our model. In real ecosystems, the fraction of direct recycling and the degradability of detritus can be controlled by the trophic structure of the food web. In aquatic ecosystems, top predators such as fish produce large quantities of highly degradable detritus (Harrault et al., 2012) that sustain a higher biomass of phytoplankton and zooplankton (Vanni and Layne, 1997; Harrault et al., 2014). In terrestrial ecosystems, herbivores also produce excrements that are easily degraded by the soil community and lead to an increase of the primary production (McNaughton, 1984; Belovsky and Slade, 2000). Primary producers can also strongly influence decomposition. In terrestrial ecosystems, plant leaf traits affect the composition and the quality of the litter (Cornwell et al., 2008). Primary producer stoichiometry is also highly variable both between and within species (Sterner et al., 2002; Dickman et al., 2006; Danger et al., 2007, 2009; Mette et al., 2011), which affects detritus quality and stoichiometry. While our results remain qualitatively robust to different values of C:N ratios of primary producers (see Fig. S3-8 in the supporting information), the links between food web structure and the degradability of detritus might strongly influence food web response to nutrient enrichment through their impact on nutrient availability. In addition, primary producer stoichiometry can be a flexible trait responding to nutrient limitation or herbivory, which can limit herbivore assimilation efficiency (Branco et al., 2018), thus affecting the energy transfer in the food chain. Including these mechanisms would thus need to be tested in new versions of our model.

### Nutrient cycling and effects of feedback loops

Though we found that nutrient cycling mostly interacts with the paradox of enrichment through an enrichment effect, we also found small effects of nutrient cycling on biomass dynamics through feedback loops from all trophic levels to mineral nutrients. These effects consist in the decrease in the temporal stability of the biomass of up to 75% of consumers in C models (with nutrient cycling) compared to SC models (without nutrient cycling but with an equivalent enrichment effect) (Fig. 5). In fact, primary producers tend to be destabilised while most of the consumers from all trophic levels are stabilised (see Fig. S2-5 in the supporting information). In addition, more species have their biomass dynamics stabilised by nutrient cycling if the fraction of direct recycling *δ* is high and external inputs *I* are low. In fact, indirect recycling keeps nutrient unavailable for primary producers and tends to smooth nutrient cycling dynamics (see Fig. S2-1B and S2-1D in the supporting information). In contrast, direct recycling shortens feedback loops and then increases the coupling between each trophic levels and mineral nutrients. Such a coupling can be seen in the increased biomass CV difference between the C and SC models (see Fig. S2-6 in the supporting information) and in the increase of the total quantity of recycled nutrient CV due to the larger contribution of species direct recycling that have high CV. To try to understand better the responses of biomass dynamics to nutrient cycling feedback loops we built a food chain model (with the same parametrisation as in our food web model) to track the dynamics of each element of the system. Even if the response of biomass repartition (see Fig. S2-8 in the supporting information) and quantity of recycled nutrient (see Fig. S2-9 in the supporting information) to nutrient enrichment *I*, decomposition rate *d* and fraction of direct recycling *δ* are similar to the food web model, the effects of feedback loops on dynamics are different. The response of each trophic level depends on food chain length but the case with four species is more representative of our food web model where the maximum trophic level is equal to four for intermediate nutrient input *I* (see Fig. S2-2B and S2-2C). Unlike in our food web model, primary producers are stabilised by feedback loops while all consumer are destabilised (see Fig. S2-11C in the supporting information). Thus, mechanisms acting in food chain models, such as the predator increasing resource uptake by prey (Brown et al., 2004a), which in turn boosts primary production and reduces the unbalance between species growth rates and loss rates (Rip and McCann, 2011), do not seem to be involved in our food web model.

One important difference between the food chain and the food web models correspond to the biomass CV difference between the SC and C cases, which is actually larger in the food chain model than in the food web model. In the food web model, the CV of the total quantity of recycled nutrients is smaller by roughly one order of magnitude compared to the average species biomass CV. Nutrient cycling is the outcome of the aggregated nutrient loss from numerous species whose dynamics are not synchronous, which leads to compensation effects: when the biomasses of some species decrease, the biomasses of other species likely increase, thus keeping the total biomass and the total quantity of recycled nutrients less variable (see Fig. S2-1 and S2-7 in the supporting information). This effect is strengthened by the detritus compartment that mixes all detritus released by species and releases them at a fixed rate *d*, thus explaining the lower CV of the quantity of recycled nutrients at low fraction of direct recycling *δ*. As nutrient cycling appears to be relatively constant over time compared to species biomass dynamics, mimicking it with a constant nutrient input *I*_*recy*_ as in SC food webs leads to dynamics similar to those of the C food webs. Theory predicts that species diversity stabilises aggregated ecosystem properties through asynchronous species dynamics (Doak et al., 1998; Gonzalez and Loreau, 2008; Loreau and de Mazancourt, 2013). This rationale is supported by numerous experimental studies showing that aggregated ecosystem processes, such as primary production (Tilman, 1996; Tilman et al., 2006; Schläpfer and Schmid, 1999; Loreau, 2000; Hooper et al., 2005) or dead biomass decomposition (Knops et al., 2001; Keith et al., 2008; Gessner et al., 2010; Nielsen et al., 2011) are more stable over time than individual species dynamics and that this stability increases with the number of species. While the asynchrony linked to biodiversity seems to deeply impact the effect of nutrient cycling on food web dynamics, nutrient cycling does not affect synchrony between the biomass dynamics of species. In fact, the asynchrony between primary producer species and consumer species is not significantly modified by the presence of feedback loops (see Fig. S2-7B in the supporting information) in the food web model. Overall, our results suggest that simplicity emerges from food web dynamics, making the prediction of the impact of nutrient cycling on ecosystem functioning easier in complex food webs than in food chains. Barbier et al. (2018) found that food web properties such as biomass distribution among species can be predicted thanks to the statistical distribution of species physiological and ecological parameters. From their results, adding the statistical distribution of recycling parameters (*δ* is fixed in our study but it must vary between species) would enable us to evaluate the quantity of recycled nutrients *I*_*recy*_ and thus to predict ecosystem functioning just by knowing the overall characteristics of the community living in the ecosystem.

To sum up, our food chain and food web models respond differently to the presence of nutrient cycling loops, making the understanding of the underlying mechanisms difficult. Therefore, new models based on simple food chains and manipulating both food chain length and horizontal diversity are needed to fully understand the effects of nutrient cycling on dynamics. More generally, our results also suggest that positive effects of biodiversity on ecosystem stability might also occur through nutrient cycling. To the best of our knowledge, this hypothesis has never been fully tested in biodiversity experiments and could lead to a new research avenue. Moreover, new studies based on stochastic perturbations as in Shanafelt and Loreau (2018) would bring knowledge on the effects of nutrient cycling on other components of the stability of food chains and food webs. While previous studies suggesting that feedback loops generated by nutrient cycling are destabilising (DeAngelis, 1980), our preliminary results from our food chain model suggest that nutrient cycling can have stabilising or destabilising effects on species biomass dynamics depending on trophic levels and food chain length for instance. This discrepancy likely arises from these former results being based on a different stability measure (*i.e.* resilience instead of temporal variability) and because previous studies did not separate enrichment effects from feedback loop effects of nutrient cycling.

## Conclusion and perspectives

We identified two distinct effects of nutrient cycling. First, an enrichment effect due to the recycled nutrients that increase species persistence at low nutrient inputs by increasing resource availability but leads to a decrease in species persistence through a paradox of enrichment at higher nutrient inputs. Second, feedback loops that link each trophic level to the mineral resource through nutrient cycling increase primary producer biomass CV and decrease consumer biomass CV compared to food webs with similar nutrient availability but without recycling. However, this effect is weak in complex food webs where the effect of nutrient cycling mainly consists in an nutrient enrichment. Thus, ecologists should consider nutrient cycling in theoretical and empirical work to better predict food web response to nutrient inputs as nutrient cycling deeply changes the overall nutrient availability.

Real ecosystems are known to differ by their dependence on external inputs of mineral nutrients (Polis et al., 1997; Vadeboncoeur et al., 2003; Jickells, 2005; Bokhorst et al., 2007), and ecosystems relying less on such inputs likely depend more on nutrient cycling than ecosystems depending more on external inputs. Therefore, nutrient cycling, as suggested by our results, could influence the food webs of these ecosystems in contrasted ways. For example, in ecosystems such as eutrophic lakes (Vadeboncoeur et al., 2003) with high inputs of nutrients, nutrient cycling could mostly have a general negative effect by promoting species extinction while it could have a positive effect in ecosystems with low inputs of nutrients such as Antarctic terrestrial ecosystems (Bokhorst et al., 2007) or infertile landscapes (Hopper, 2009). In the same vein, in ecosystems with efficient nutrient cycling, nutrient losses are low so that nutrient cycling represents a very important source of nutrient and more likely leads to negative effects if the ecosystem already receive abundant external nutrient inputs.

Experiments designed to test the effects of the mechanisms involved in our model would be interesting. For example, it would be possible in mesocosms to manipulate both inputs of mineral nutrients and the efficiency of nutrient cycling (Harrault et al., 2014), *e.g.* exporting an increasing proportion of detritus, and to measure the response in terms of food web functioning and population dynamics. It would also be interesting to compare food webs of different types of natural ecosystem with contrasting nutrient cycling and mineralisation rates. Typically, our model probably better corresponds to an aquatic food web (*i.e.* fully size-structured web) and aquatic and terrestrial food webs should be compared.

Even if the detritus compartment affects the effects of nutrient cycling on food webs, its role cannot be fully appreciated in our model because there are no decomposers and no brown food web. In fact, detritus are more than a transient pool for nutrients since, in real food webs, they are resources for decomposers and are recycled through the whole brown food web (Moore et al., 2004). Another important step will be to include in models a true brown food web containing decomposers feeding on detritus in parallel to the green food webs relying on photosynthesis (Moore et al., 2004; Zou et al., 2016). The interactions between green and brown food webs are conditioned by stoichiometric constrains on primary producers and detritus that affect the competition/mutualist interactions between primary producers and decomposers (Daufresne and Loreau, 2001; Cherif and Loreau, 2013; Zou et al., 2016)

To go further, the flexible stoichiometry of primary producers (and phytoplankton in particular) can also deeply affect food web dynamics and consumer persistence as it can limit herbivore assimilation efficiency (Loladze et al., 2000; Branco et al., 2018). In fact, Urabe and Sterner (1996) demonstrated experimentally that increasing light availability first increases phytoplankton and zooplankton biomass productions but then led to zooplankton extinction because of the low nutritional quality of phytoplankton biomass if the light to nutrient ratio was to high.

Thus, stoichiometric constraints and green and brown food web interactions can deeply change the functioning and the stability of ecosystems (Daufresne and Loreau, 2001; Moore et al., 2005; Attayde and Ripa, 2008; Zou et al., 2016) but these results have so far not been tested in complex food web models.

## Supporting information

Supporting information final

## Acknowledgement

We thank the École Normale Supérieure, the PhD program "École Doctorale Frontières du Vivant (FdV) – Programme Bettencourt" and the Initiative Structurante Ecosphère Continentale et Côtière (IS EC2CO) for their financial support. The simulations were performed at the HPCaVe at UPMC-Sorbonne Université. We also thank the anonymous reviewers, Shawn Leroux, Jean-François Arnoldi, Wojciech Uszko and Samraat Pawar for their detailed reviews that strongly helped us to improve this manuscript. Version 7 of this preprint has been peer-reviewed and recommended by Peer Community In Ecology (doi:10.24072/pci.ecology.100046).

## Supporting information, data and code accessibility

The supporting information (see Supporting_information_final.pdf) contains the following section:

- S1 Appendix – Parameter calculation
- S2 Appendix – Complementary results
- S3 Appendix – Sensitivity analysis

The C++ code of the simulations and the R code of the figures are available on Zenodo (doi:10.5281/zenodo.3697083).

