## Supporting information final for "Interplay between the paradox of enrichment and nutrient cycling in food webs"

Peer Community In
Ecology

### Interplay between the paradox of enrichment and nutrient cycling in food webs

Pierre Quévreux[1], Sébastien Barot[1], & Élisa Thébault[1]

[1] Sorbonne Université, Sorbonne Paris Cité, Paris Diderot Univ Paris 07, CNRS, INRA, IRD, UPEC, Institut d'Écologie et des Sciences de l'Environnement – Paris, iEES-Paris, 4 place Jussieu, F-75252 Paris, France

#### S1 Appendix - Parameter calculation

##### Allometric parameter calculation

The value of the primary producers growth rate was taken from Savage et al. (2004) and Binzer et al. (2012):

$$r_i = e^{I_r} M_i^{-1/4} e^{Ea_r(T_0 - T/kTT_0)} \tag{11}$$

| | |
|---|---|
| $e^{I_r}$ | allometric scaling constant at 20°C ($g^{1/4}.s^{-1}$) |
| $M_i$ | body mass ($g$) |
| $e^{Ea_r(T_0 - T/kTT_0)}$ | temperature dependency term |

We considered that temperature was constant at 20°C (thus $T = T_0$) and with $I_r = -15.68$ (Binzer et al., 2012) we have:

$$r_i = r M_i^{-1/4} \tag{12}$$

With $r = 0.87\,kg^{1/4}.year^{-1}$. Metabolic rates were taken from Brose et al. (2006b) and Brose (2008) with $x/r = 0.138$ for primary producers, $x/r = 0.314$ for invertebrates and $x/r = 0.88$ for ectotherm vertebrates. Since we did not apply the time scale normalisation by the growth rate of primary producers as done in Brose et al. (2006b), we have $x = 0.12$ for primary producers, $x = 0.27$ for invertebrates and $x = 0.78$ for ectotherm vertebrates. We used the values for invertebrates for consumers in our simulations.

##### Handling time

In this model, the handling time $h_{ij}$ also follows an allometric scaling. We used the expression defined by Petchey et al. (2008) and also used by Thierry et al. (2011). The original expression has been divided by prey body mass to have a mass specific allometric parametrisation:

$$h_{ij} = \begin{cases} \dfrac{h_i}{b - \dfrac{M_j}{M_i}} \dfrac{1}{M_j} & \text{if } \dfrac{M_j}{M_i} < b \\ \infty & \text{if } \dfrac{M_j}{M_i} > b \end{cases} \tag{13}$$

| | |
|---|---|
| $h_i$ | allometric scaling constant ($year.kg^{-1}$) |
| $b$ | maximum predator-prey body mass ratio ($0.05$) |
| $M_i$ | body mass of the predator ($kg$) |
| $M_j$ | body mass of the prey ($kg$) |

The maximum prey-predator body mass ratio $b$ delimits the diet breadth. The handling time function is U-shaped (Fig. S1-1) if the predator-prey body mass ratio is below $b$, otherwise the handling time tends to infinity and the prey is not consumed by the predator. Unfortunately, no values of the allometric scaling constant $h_i$ could be found in the literature. However, the maximum ingestion rate $y_i$ is well quantified (Yodzis and Innes, 1992; Brose et al., 2006b; Vucic-Pestic et al., 2010) and corresponds to the reverse of the handling time. Following Brose et al. (2006b), the ingestion rate is set proportional to the metabolic rate:

$$y_i = yx_i \tag{14}$$

With $y = 8$ for invertebrates and $y = 4$ for ectotherm vertebrates. Then, we assumed that the values from Brose et al. (2006b), which do not depend on the body mass of the prey, are the average over all possible prey body masses (interval $[0, bM_i]$ defined in equation 13). Thus, we can state that:

$$y_i = \frac{1}{bM_i} \int_0^{bM_i} \frac{1}{h_{ij}} dM_j \tag{15}$$

Thus, by replacing $h_{ij}$ by the expression from equation 13:

$$\begin{aligned} y_i &= \frac{1}{bM_i} \int_0^{bM_i} \frac{1}{\dfrac{h_i}{b - \dfrac{M_j}{M_i}} \dfrac{1}{M_j}} dM_j \\ &= \frac{1}{h_i bM_i} \int_0^{bM_i} (b - \frac{M_j}{M_i}) M_j dM_j \\ &= \frac{1}{h_i bM_i} \left[ \frac{bM_j^2}{2} - \frac{M_j^3}{3M_i} \right]_0^{bM_i} \\ &= \frac{b^2 M_i}{6h_i} \end{aligned} \tag{16}$$

Thus:

$$h_i = b^2 M_i / 6y_i \tag{17}$$

And by replacing $h_i$ in equation 13 by the expression found in equation 17:

$$h_{ij} = \frac{b^2}{6y_i(b - \dfrac{M_j}{M_i})} \frac{M_i}{M_j} \tag{18}$$

$y_i$ is defined as in equation 4a:

$$y_i = yM_i^{-1/4} \tag{19}$$

| | |
|---|---|
| $y$ | allometric scaling constant ($kg^{1/4}.year^{-1}$) expressed as $8.x$ (Brose et al., 2006b) |
| $M_i$ | body mass of the organism ($kg$) |

Handling time is minimum for $M_j = \frac{b}{2}M_i$. The value of the maximum predator-prey body mass ratio $b$ is set to 0.05 so that the handling time is minimal for prey 40 times smaller than their predators. This value is consistent with the average predator-prey body mass ratio found by Brose et al. (2006a). To limit the number of equations, the interactions involving prey out of the interval $[0.1bM_i, bM_i]$ were neglected.

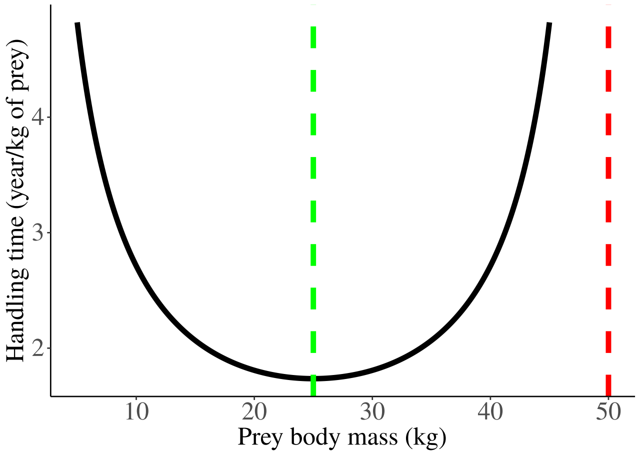


**Figure S1-1.** Handling time as a function of prey body mass ($b = 0.05$, $M_i = 1000kg$). The red dashed line represents the upper limit of prey body mass that predators can handle ($M_i/M_j < b$) and the green dashed line represents the optimal prey body mass minimising the handling time ($M_j = \frac{b}{2}M_i$).

#### Stoichiometry and C:N ratios

The limiting nutrients considered in our model could be any mineral nutrient but we chose nitrogen to parametrise the carbon to nutrient ratio. C:N ratios were taken from data of pelagic communities (Anderson, 1992) with C:N=6.6 for primary producers (value for phytoplankton) and C:N=5 for consumers (average C:N ratios of bacteria (5.1), protozoa (5.5) and copepods (4.67)). The amount of nutrients released by consumers from non-assimilated prey biomass depends on both the C:N ratio of prey and consumers. The C:N ratio of non-assimilated biomass $\alpha_{Dij}$ can be calculated by using the constraints on mass conservation and maintenance of species homoeostasis (equation 9). The ingested biomass by consumer species $i$ of prey species $j$ contains a mass $C_j$ of carbon and $N_j$ of nutrients ($\alpha_j = C_j/N_j$). A fraction $e_{ij}$ of $C_j$ is converted into a mass $C_i$ of carbon of the consumer while the remaining fraction $1 - e_{ij}$ is converted into a mass $C_{Dij}$ of detritus. We define $N_i$ as the assimilated mass of nutrients by the consumer ($\alpha_i = C_i/N_i$) and $N_{Dij}$ as the non assimilated mass of nutrients excreted in the detritus pool (with $\alpha_{Dij} = C_{Dij}/N_{Dij}$). By mass conservation, we have the two following relations:

$$N_j = N_i + N_{Dij} \tag{20a}$$

$$C_j = C_i + C_{Dij} = e_{ij}C_j + (1 - e_{ij})C_j \tag{20b}$$

As the predator and the prey keep their C:N ratios $\alpha_i$ and $\alpha_j$ constant, we can derive the expression of $\alpha_{Dij}$ as a function of $e_{ij}$, $\alpha_i$ and $\alpha_j$:

$$\begin{aligned} \frac{C_j}{\alpha_j} &= \frac{C_i}{\alpha_i} + \frac{C_{Dij}}{\alpha_{Dij}} = \frac{e_{ij}C_j}{\alpha_i} + \frac{(1 - e_{ij})C_j}{\alpha_{Dij}} \\ \alpha_{Dij} &= \frac{\alpha_j\alpha_i(1 - e_{ij})}{\alpha_i - \alpha_j e_{ij}} \end{aligned} \tag{21}$$

#### Adaptive foraging equation

We detail in this part the expression of $\partial g_i / \partial \omega_{ik}$ that is part of equation (6). $g_i$ is the total growth rate of species $i$ and is defined as $\frac{dB_i}{dt} = g_i B_i$. However, we notice that :

$$\frac{\partial g_i}{\partial \omega_{ik}} = \frac{\partial}{\partial \omega_{ik}} \left( \sum_{\ell = prey} e_{i\ell} F_{i\ell} \right) \tag{22}$$

As only $F_{i\ell}$ depends on $\omega_{ik}$ in equation (3b). Thus:

$$\begin{aligned}
\frac{\partial g_i}{\partial \omega_{ik}} &= e_{ik} \frac{\partial F_{ik}}{\partial \omega_{ik}} + \sum_{\ell \neq k} e_{i\ell} \frac{\partial F_{i\ell}}{\partial \omega_{ik}} \\
&= e_{ik} \left( \frac{a_i B_k (1 + \sum_m \omega_{im} a_i h_{im} B_m) - \omega_{ik} a_i B_k (a_i h_{ik} B_k)}{(1 + \sum_m \omega_{im} a_i h_{im} B_m)^2} \right) \\
&\quad + \sum_{\ell \neq k} \frac{e_{i\ell} \omega_{i\ell} a_i B_\ell (-a_i h_{ik} B_k)}{(1 + \sum_m \omega_{im} a_i h_{im} B_m)^2} \\
&= \frac{e_{ik} a_i B_k + \sum_m e_{ik} a_i^2 \omega_{im} h_{im} B_k B_m - e_{ik} a_i^2 \omega_{ik} h_{ik} B_k^2 - \sum_{\ell \neq k} e_{i\ell} a_i^2 \omega_{i\ell} h_{ik} B_k B_\ell}{(1 + \sum_m \omega_{im} a_i h_{im} B_m)^2} \\
&= \frac{e_{ik} a_i B_k + \sum_{m \neq k} e_{ik} a_i^2 \omega_{im} h_{im} B_k B_m - \sum_{\ell \neq k} e_{i\ell} a_i^2 \omega_{i\ell} h_{ik} B_k B_\ell}{(1 + \sum_m \omega_{im} a_i h_{im} B_m)^2} \\
&= \frac{a_i B_k \left( e_{ik} + \sum_{\ell \neq k} a_i \omega_{i\ell} B_\ell (e_{ik} h_{i\ell} - e_{i\ell} h_{ik}) \right)}{(1 + \sum_m \omega_{im} a_i h_{im} B_m)^2}
\end{aligned} \tag{23}$$

### S2 Appendix - Complementary results

#### Complex food webs

##### Overview of the dynamics of the food web

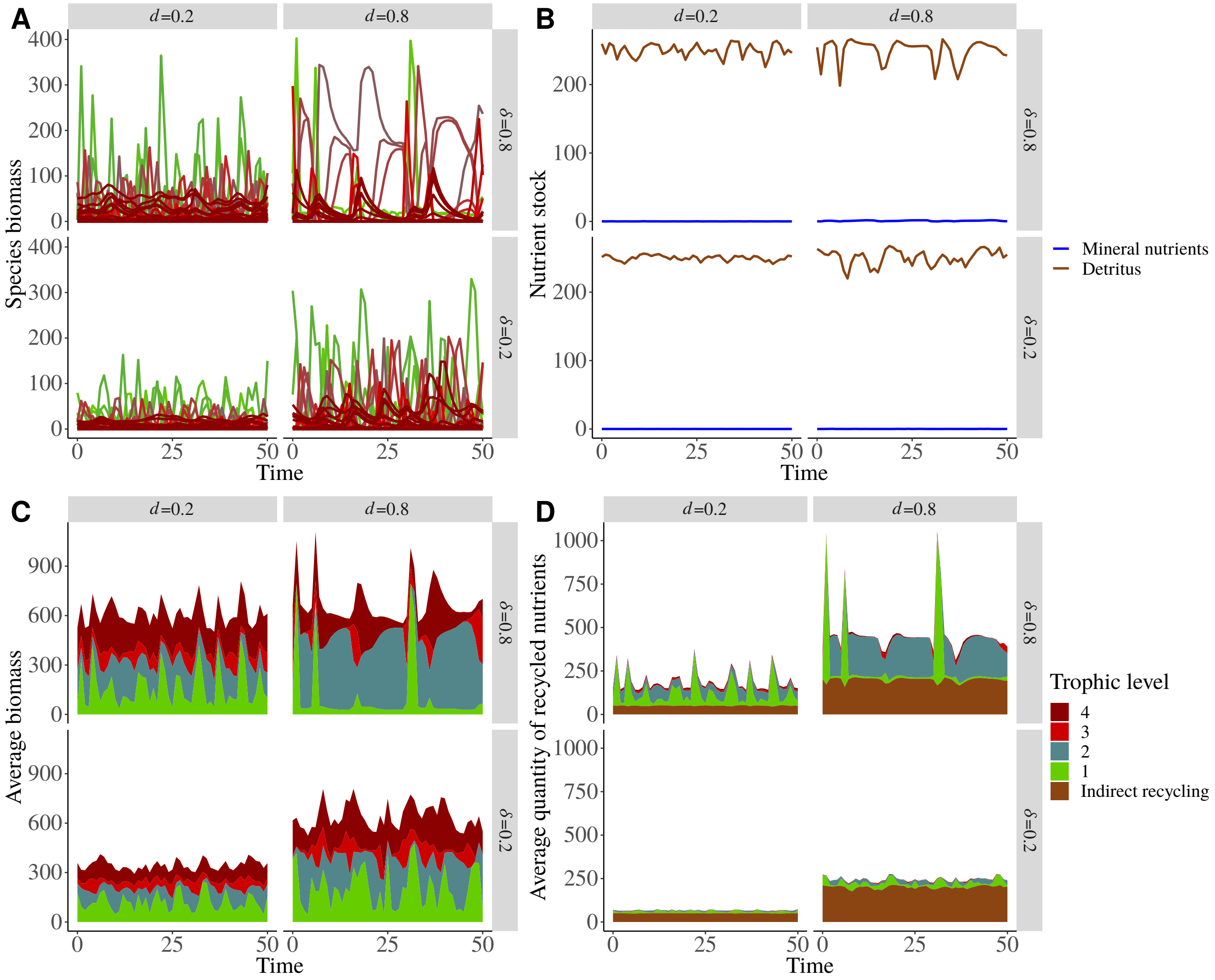


**Figure S2-1.** Dynamics of species biomasses, abiotic compartments and nutrients recycled in a C food web for $I = 50$. **A)** Biomass dynamics of each species. Lower trophic levels are in green and higher trophic levels are in dark red. **B)** Mineral nutrients (blue) and detritus (brown) dynamics. **C)** Species biomasses aggregated by trophic levels (from green to dark red areas). **D)** Nutrients directly recycled by species aggregated by trophic levels and indirectly recycled nutrients (brown area).

Our complex food web model generates highly variable species biomasses (Fig. S2-1A) while aggregated biomasses are relatively less variable (Fig. S2-1C). The quantity of recycled nutrients by each species is also highly variable while the total quantity of recycled nutrients is less variable (Fig. S2-1D). In addition, the aggregated quantities of recycled nutrients are more stable than the quantities of recycled nutrients at species level. We also observe that primary producers and herbivores are the main contributors to nutrient cycling (Fig. S2-1D) respectively because of their high biomass and their low assimilation efficiency ($e_{ij} = 0.45$). The low contribution of the carnivores can be attributed to their high assimilation efficiency ($e_{ij} = 0.85$) and their low metabolic rate due to their large body mass.

#### Complementary results on species dynamics

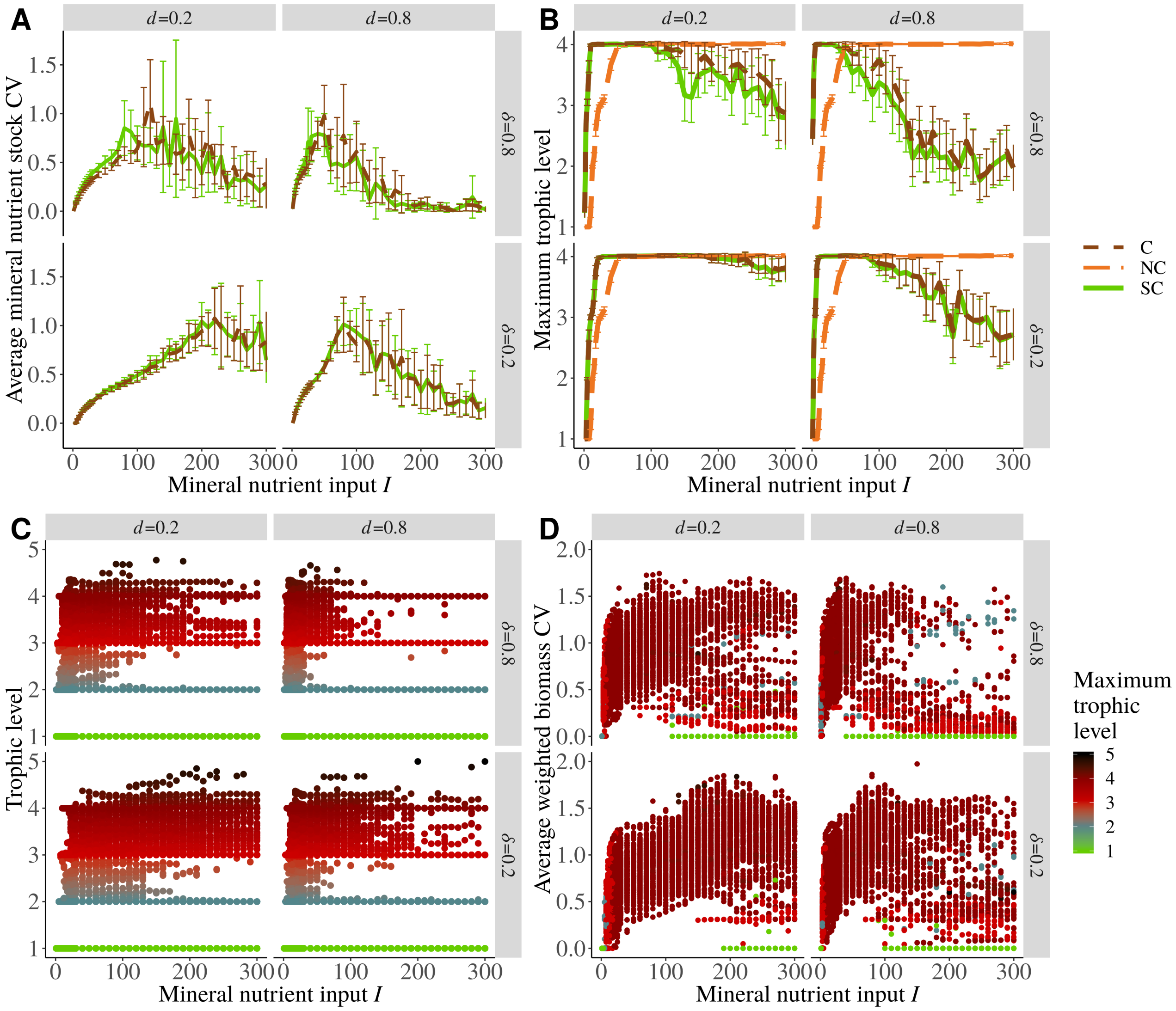


**Figure S2-2.** Overall response of food webs to nutrient cycling and to an enrichment gradient $I$ as a function of recycling parameters $d$ (decomposition rate) and $\delta$ (fraction of direct recycling). **A)** Average coefficient of variation of the mineral nutrient stock. The C food webs (dashed brown) are the food webs with nutrient cycling and the SC food webs (solid green) are the food webs without nutrient cycling but with a simulated enrichment due to nutrient cycling. **B)** Average maximum trophic level. The trophic level 1 corresponds to primary producers and NC food webs (long-dashed orange) are the food webs without nutrient cycling. **C)** Trophic level of each species in each simulated C food web. The colour gradient also represents the trophic levels (from primary producers in green to top predators in black). **D)** Average species biomass CV in each simulated C food web. The colour scale represents the maximum trophic level sustained by each simulated food web.

Nutrient stock CV (Fig. S2-2A) responds similarly to nutrient enrichment and nutrient cycling feedback loop presence than the CV of species biomass (Fig. 4A). Indeed, we first see an increase followed by a decrease with increasing mineral nutrient inputs $I$. In addition, we do not see significant differences between C food webs with nutrient cycling and SC food webs without nutrient cycling but with a simulated enrichment due to nutrient cycling, thus, nutrient cycling does not seem to modify the general variability of the mineral nutrient stock.

In food webs with nutrient cycling, maximum trophic level (Fig. S2-2B) follows a hump-shaped relationship with external nutrient inputs: first there is a sharp increase in food web maximum trophic level for low nutrient inputs, then a plateau and finally a decrease in food web maximum trophic level for high nutrient inputs. The decrease of the maximum trophic levels is correlated to the decrease of persistence, suggesting that higher trophic levels are the first species that get extinct due to the paradox of enrichment.

Fig. S2-2C describes the general distribution of trophic levels in simulated food webs. Lower trophic levels tend to be

well separated with consumer eating prey one trophic level below. The similar body masses of primary producers added to the constraint of the feeding niche must lead to this structure but we also notice that omnivory occurs more frequently in higher trophic levels.
The average CV of species biomass in a food web is correlated with the maximum trophic level of the food web (Fig. S2-2D). It is high when food webs have at least two trophic levels (and seems to be higher if high trophic levels persist), or it is null when food webs contain only primary producers as the system reaches fix points.

**Response of nutrient stocks and flows to nutrient enrichment**

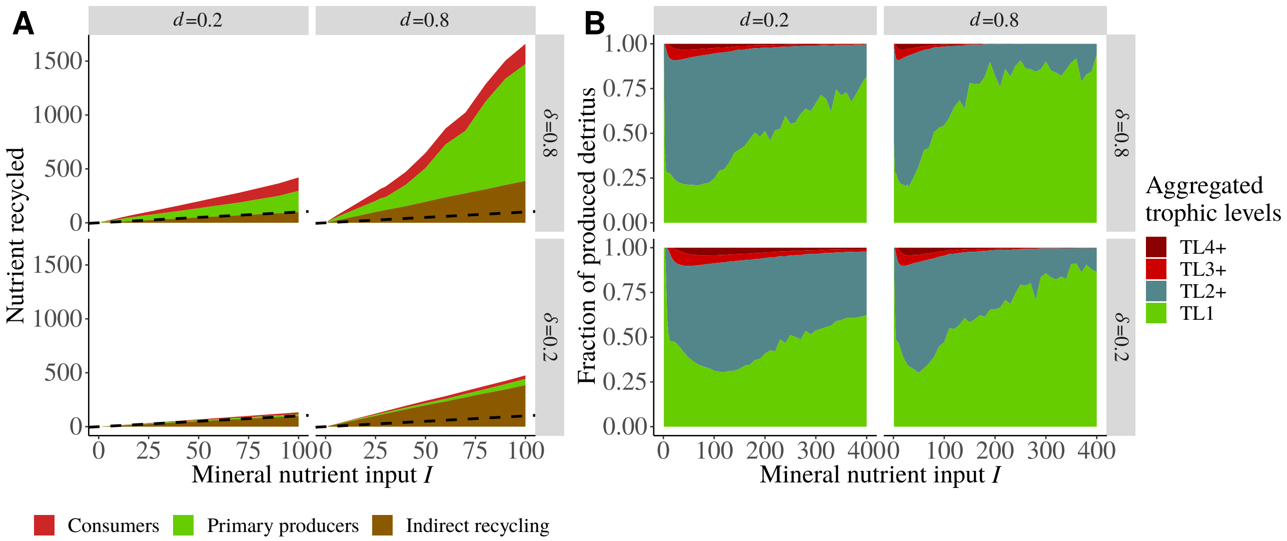


**Figure S2-3.** Detailed origins of recycled nutrients and detritus. **A)** Zoom in Fig. 3A for $I \in [0, 100]$. **B)** Fraction of detritus produced by each aggregated trophic level (fraction $(1-\delta)$ of indirect recycling plus fraction $(1-e_{ij})$ of non assimilated biomass, see equation (7b)). TL1 represents basal species (primary producers) and higher trophic levels are aggregated. Species with a trophic level comprised in the interval $[i, i+1[$ are in the group TLi+. For instance, TL2+ gathers strict herbivores and omnivores eating both primary producers and herbivores.

As explained in the main text, the quantity of recycled nutrients increases with external nutrient inputs $I$, the fraction of direct recycling $\delta$ and the decomposition rate $d$ (Fig. S2-3A). In detail, primary producer and consumer contributions vary with $I$. At low nutrient inputs $I$, consumers contribute significantly to nutrient recycling as they have a high biomass (Fig. S2-4A). This must be explained by the high species persistence (Fig. 3B in the main text) combined with high trophic levels survival (Fig. S2-2B). Moreover, herbivores contribute strongly (Fig. S2-3B) to indirect recycling due to their low assimilation efficiency ($e_i = 0.45$) that release a lot of detritus when they consume primary producer biomass. However, at high external inputs $I$, primary producers are responsible of most of direct (Fig. S2-3A) and indirect (Fig. S2-3B) recycling, due to their sheer biomass that is much higher than consumer biomass (Fig. S2-4A).

Mineral nutrient stock is negligible compared to detritus stock that increases linearly with external nutrient inputs (Fig. S2-4D), suggesting that it is controlled by primary producers in the food web. As the external inputs are balanced by the losses from the mineral nutrient and detritus compartments ($I = \ell N + \ell D$) (see equations (7a) and (7b)), detritus loss must balance the quasi totality of external nutrient inputs, leading to $I \simeq \ell D$. This explains the linear increase of detritus stock and their insensitivity to recycling parameters $d$ and $\delta$. While mineral nutrient and detritus stocks are not affected by $d$ and $\delta$, increasing $d$ and $\delta$ mainly increases the flows between those compartments. In fact, increasing $d$ and $\delta$ increases primary production through nutrient availability (Fig. S2-4B), that is balanced by a higher mortality due density dependent mortality (Fig. S2-4C) that quadratically increases when biomass linearly increases. This increased mortality increases the quantity of recycled nutrients (Fig. 3A) that fuels biomass production (Fig. S2-4B).

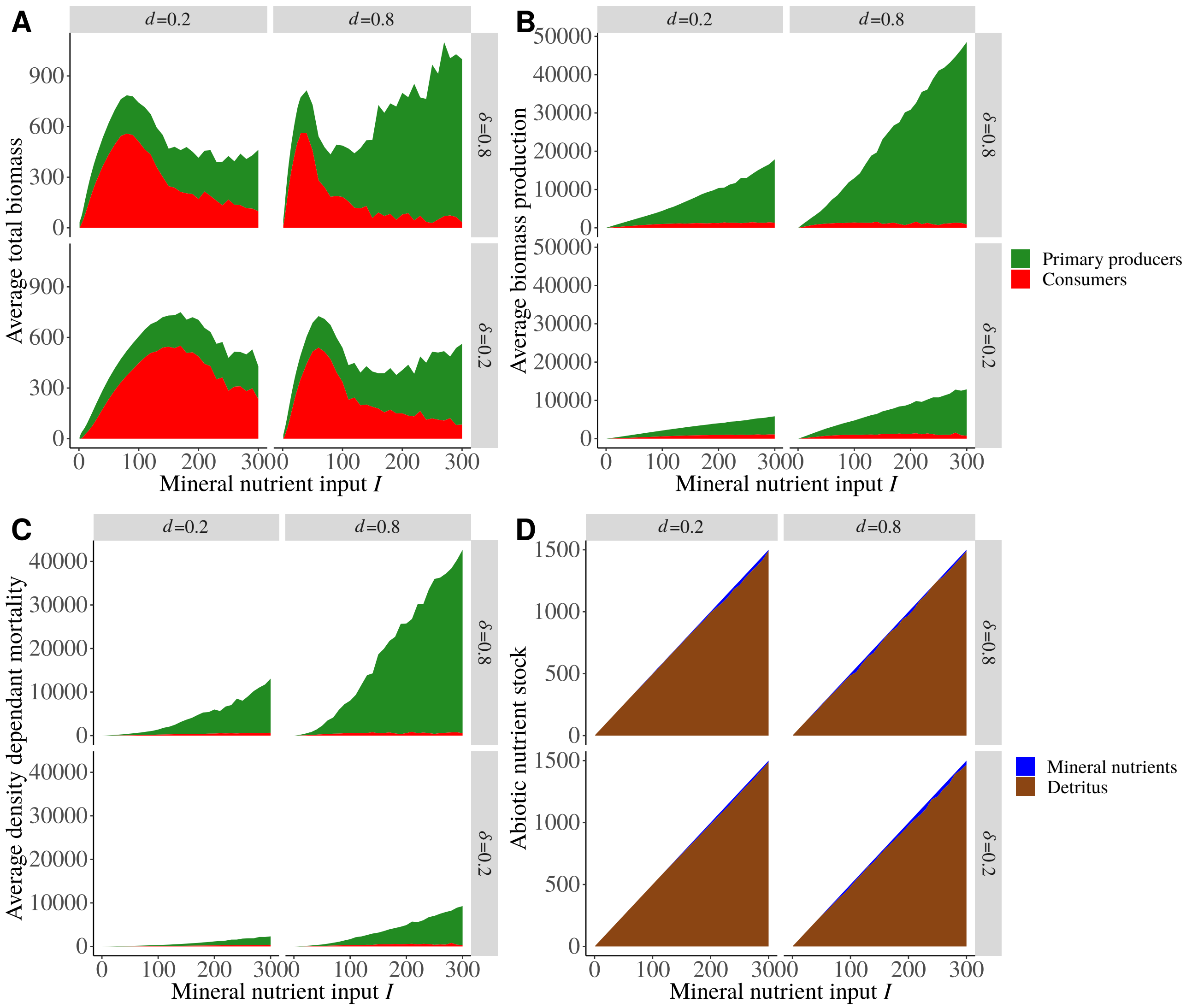


**Figure S2-4.** Response of biomass, nutrient stocks and flows to an enrichment gradient $I$ as a function of recycling parameters $d$ (decomposition rate) and $\delta$ (fraction of direct recycling). Only C food webs (with nutrient cycling) are presented. **A)** Average aggregated biomass of primary producers (green) and consumers (red). **B)** Average primary and secondary productions. 100 food webs are simulated and only values of primary producer and consumer biomass and production where these categories of species persisted are kept. **C)** Average cumulated biomass lost due to density dependent mortality ($\sum \beta_i B_i^2$). **D)** Average mineral nutrient (blue) and detritus stocks in C food webs with at least one persisting species.

##### Comparison between species biomass CV in C and SC models

The effects of the presence of recycling loops depend on the considered trophic level. Primary producers dynamics (Fig. S2-5A) are mostly destabilised by recycling feedback loops but they become largely unaffected at high nutrient inputs as they are the last surviving species (system with only primary producers reach fixed points with CV=0). Consumers, whatever their trophic level, have less variable biomasses in presence of nutrient cycling feedback loops (Fig. S2-5B-D). The response of primary producers and consumers to recycling parameters ($d$ and $\delta$) are the same than the general response presented in Fig. 5. In addition, the destabilising effect for primary producers and stabilising effects for consumers of recycling feedback loops are stronger (*i.e.* larger CV difference) when they occur for more species (for $\delta = 0.8$ in Fig. S2-6).

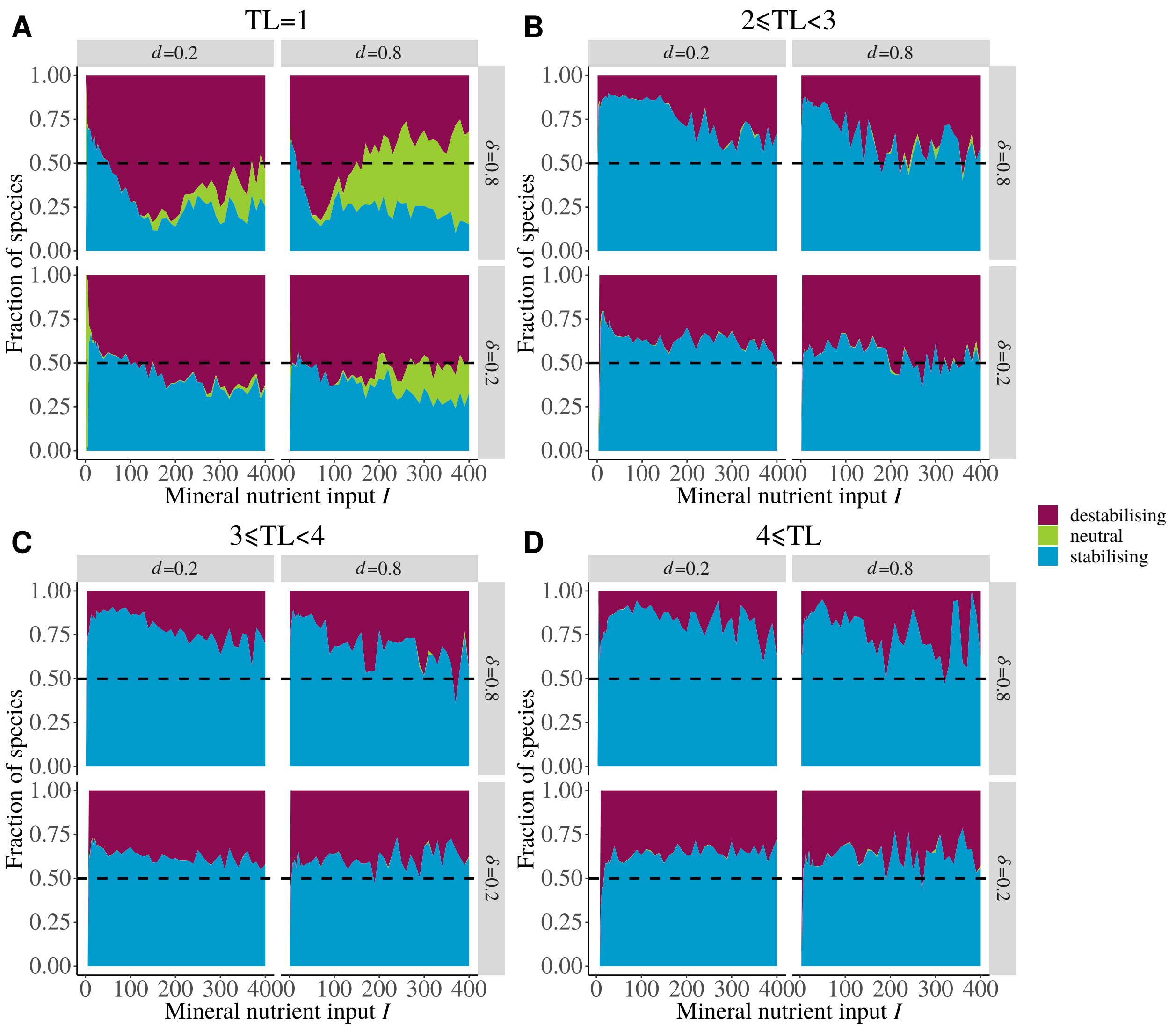


**Figure S2-5.** Effect of nutrient cycling on biomass CV at species level. For each combination of parameters, the biomass CV of the same species between C and SC food webs is compared. If the CV is higher in the SC food web (without nutrient cycling but with a mineral nutrient input simulating the enrichment effect of nutrient cycling) than in the C food web (with nutrient cycling), then nutrient cycling feedback loops have a stabilising effect on dynamics. If the difference is bellow $10^{-4}$, recycling loops are assumed to have neutral effects on dynamics. We also consider species extinction in the SC food web and not in the C food web as a stabilising effect of recycling feedback loops. The fraction of stabilised or destabilised species among all simulated food webs gives the overall effect of nutrient cycling feedback loops at species level dynamics for **A)** primary producers, **B)** herbivores and omnivorous carnivores, **C)** predators with a trophic level comprised between 3 and 4 and **D)** top predators.

The temporal stability of ecosystem aggregated processes, such as total biomass production, relies on the asynchrony between the dynamics of each species Yachi and Loreau (1999); McCann (2000); Hooper et al. (2005). Asynchrony can be calculated as the ratio of the CV of the aggregated process to the average CV from species level dynamics following Loreau and de Mazancourt (2008) (see also Fig. S2-7). If the synchrony $\varphi$ is low (its value is between 0 and 1), then the aggregated process dynamics are less variable than each of its component dynamics.

The synchrony between the quantity of nutrient directly recycled by each species (Fig. S2-7A) or between the biomasses of each species (Fig. S2-7B) first drops at very low nutrient inputs $I$ and then slowly increase with $I$. This is the opposite to the response of species persistence that first increases and then decreases with $I$ (see Fig. 3B in the main text). Therefore, synchrony must be directly linked to species persistence as species richness promotes the buffuring effect of aggregated processes (Yachi and Loreau, 1999). Finally, the recycling parameters $d$ (decomposition rate) and $\delta$ (fraction of direct recycling) have no effect on biomass synchrony. Thus, the stabilising effects seen in Fig. S2-5 and S2-6 are likely not due to changes in the asynchrony between species biomasses or recycling.

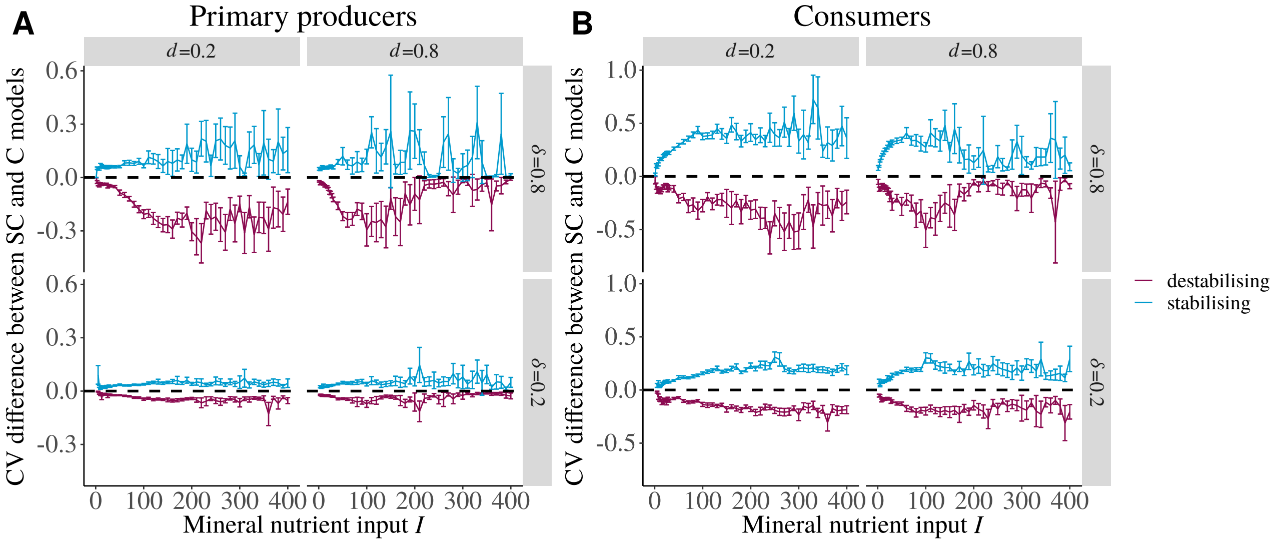


**Figure S2-6.** Average CV difference between SC models and C models in cases where species are stabilised or destabilised for **A)** primary producers and **B)** consumers.

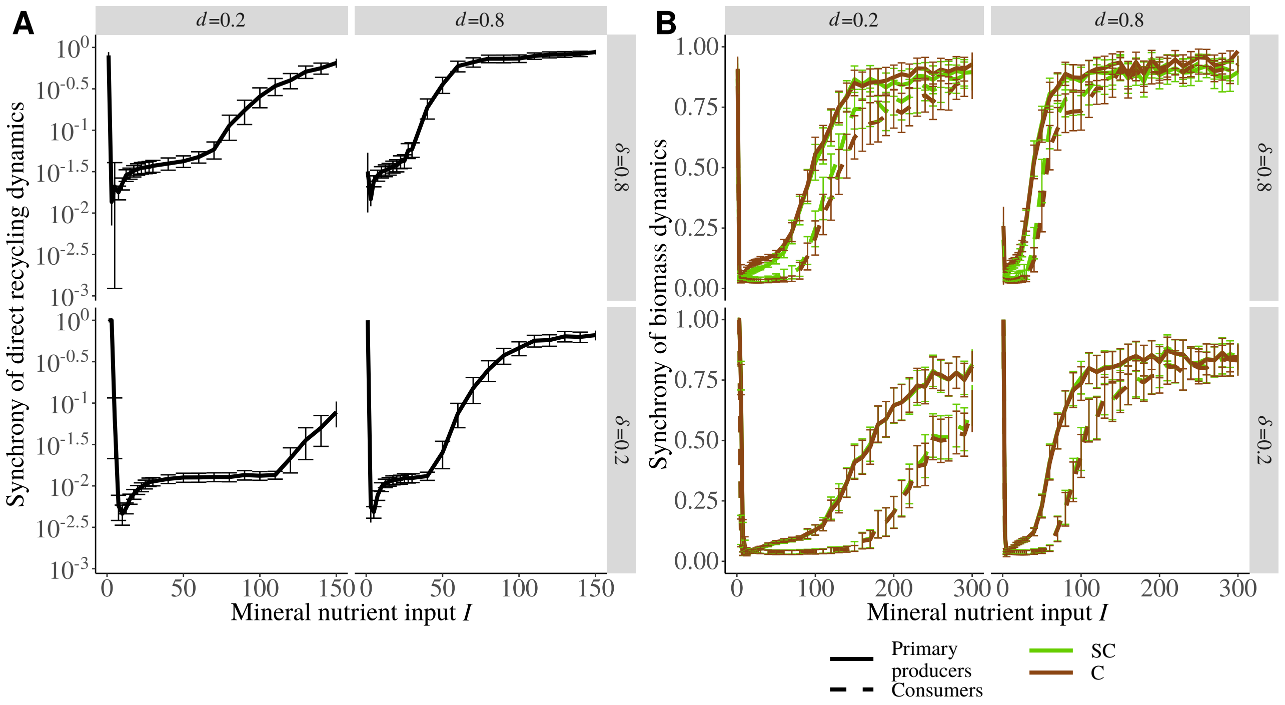


**Figure S2-7. A)** Average synchrony between the quantities of directly recycled nutrient by each species in C food webs (with nutrient cycling). It is calculated as $\varphi = CV_{I_{recy}}^2/\overline{CV}_{S_{recy}}^2$ with $CV_{I_{recy}}$ the CV of the total quantity of recycled nutrients and $\overline{CV}_{S_{recy}}$ the average CV of the quantity of nutrient directly recycled by each species weighted by the quantity of nutrient directly recycled by each species. **B)** Average synchrony between species biomass dynamics. It is calculated as $\varphi = CV_{tot}^2/\overline{CV}_S^2$ with $CV_{tot}$ the CV of the total biomass and $\overline{CV}_S$ the average species biomass CV weighted their biomass. Solid lines are the values for primary producers and dashed lines for consumers. Brown lines are the values calculated in C food webs (with nutrient cycling) and green lines values from SC food webs (without nutrient cycling but with a simulated enrichment due to nutrient cycling).

#### Food chain study

##### The food chain model

The food chain model is a simplified version of the food web model, with only four species, a primary producer, a herbivore, a carnivore and a top-predator. It is thus built with the same equations and the same parameters as the food web model except for the adaptive foraging that is not relevant in such a model. In the simulations, the body masses of the four species are respectively $10^{-4}$, $4.10{-}3$, 0.16 and 6.4 kg (each consumer being 40 times bigger than its prey), and their initial biomass are respectively 1, 0.5, 0.1 and 0.1 kg.v$^{-1}$.

##### Overall response of the food chain

The biomasses of the different species form a bottom-heavy pyramid, higher trophic levels being rare (Fig. S2-8). Adding trophic levels also changes biomass repartition: an even food chain length (Fig. S2-8A and S2-8C) leads to a herbivore biomass higher and a primary producer biomass lower than in a food chain with an odd food chain length (Fig. S2-8B). The total biomass increases with external nutrient inputs $I$ and this increase is sharper for high values of decomposition rate $d$ or fraction of direct recycling $\delta$. For $\delta = 0.8$ the food chain even collapses at $I \simeq 120$. Compared to the food chain model, consumer total biomass is much higher in the food web model (Fig. S2-4A) because there are more consumer species in the food web model than in the food chain model.

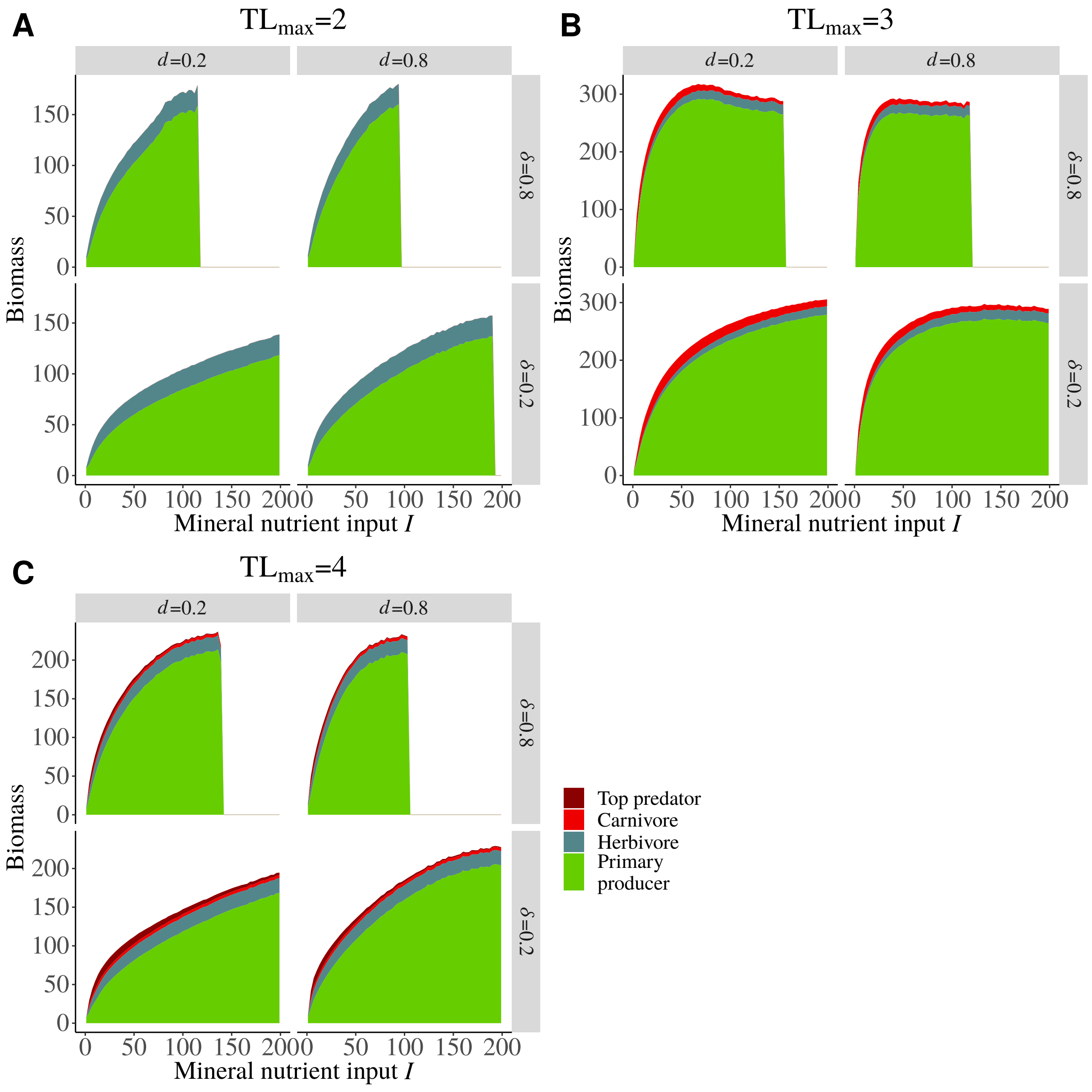


**Figure S2-8.** Average biomass of primary producers (green), herbivores (light blue), carnivores (red) and top predators (dark red) in C food chains. Three food chain lengths are tested: **A)** two species, **B)** three species and **C)** four species.

Nutrient cycling also represents a significant part of the total nutrient input in the mineral nutrient pool (Fig. S2-9) but it is slightly less important than in the food web model probably due to the lower total biomass in the food chain model. Increasing the fraction of direct recycling $\delta$ and the decomposition rate $d$ increase the quantity of recycled nutrientsas found in the food web model. The total quantity of recycled nutrients is also sensitive to food chain length. Food chains with even food chain length (Fig. S2-9A and S2-9C) recycle more nutrients than food chains with odd food chain length (Fig. S2-9B). Total biomass has exactly the same response (Fig. S2-8) because of trophic cascades: with even food chain length, primary producers are controlled by herbivores and as they are the most abundant species, the total biomass decrease. As nutrient cycling directly depends on species biomass (see equations (3a) and (3b)), the quantity of recycled nutrients also follows the trophic cascade pattern.

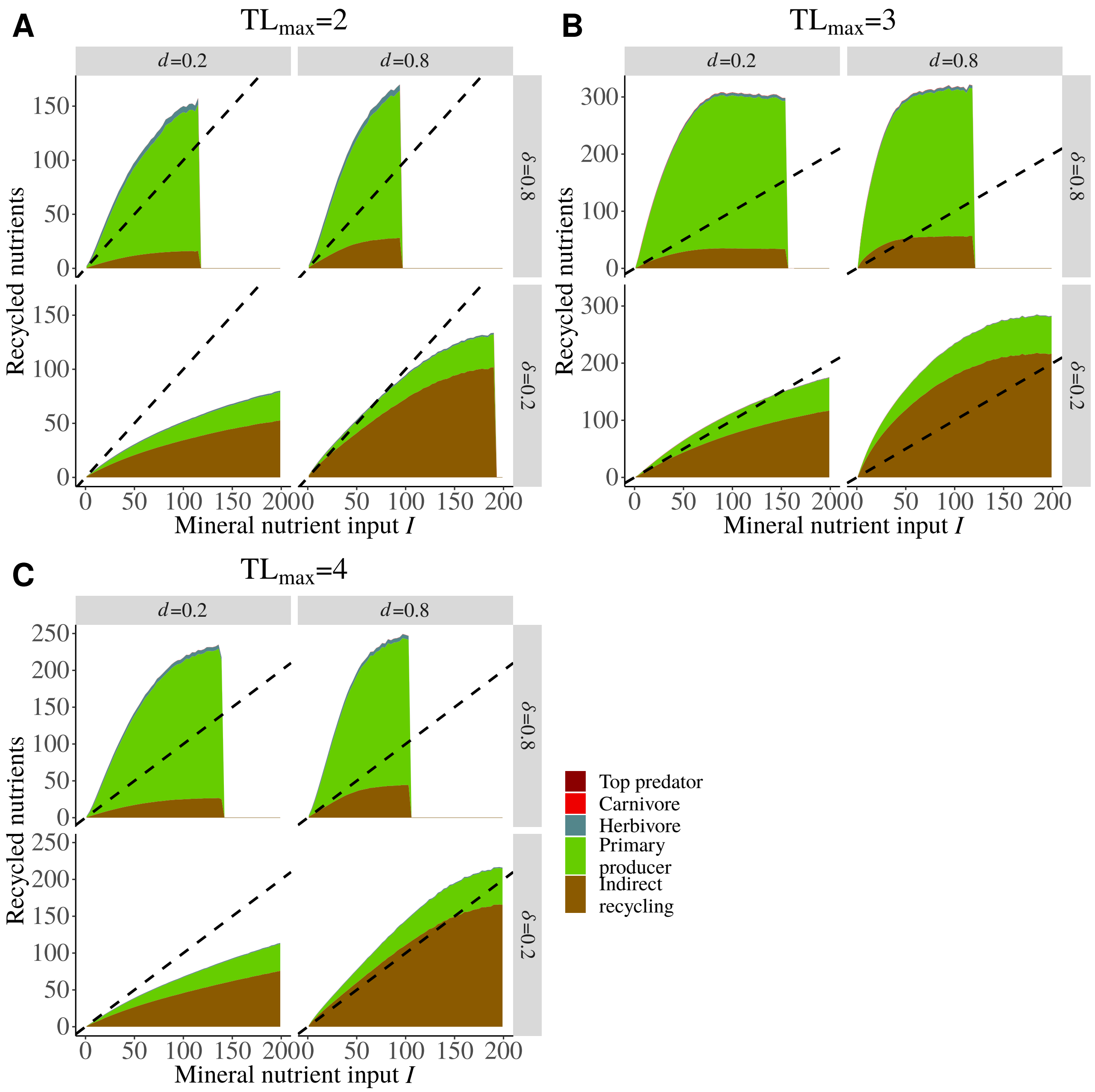


**Figure S2-9.** Average quantity of nutrients directly recycled by primary producers (green), herbivores (light blue), carnivores (red) and top predators (dark red) and indirectly (brown) in C food chains. Three food chain lengths are tested: **A)** two species, **B)** three species and **C)** four species.

Species biomass and detritus stock CV increase with external nutrient input $I$ (Fig. S2-10). However, primary producer biomass in food chains with even food chain length only (Fig. S2-10A and S2-10C) first increase and then decrease with $I$. Mineral nutrient stock CV has the same relation with $I$ whatever the food chain length. Such an increase of biomass CV is consistent with our results from the food web model (Fig. 4A in the main text) and the paradox of enrichment predictions, except for primary producers whose biomass CV decreases with $I$. This result is counter-intuitive and at this point we cannot explain it.

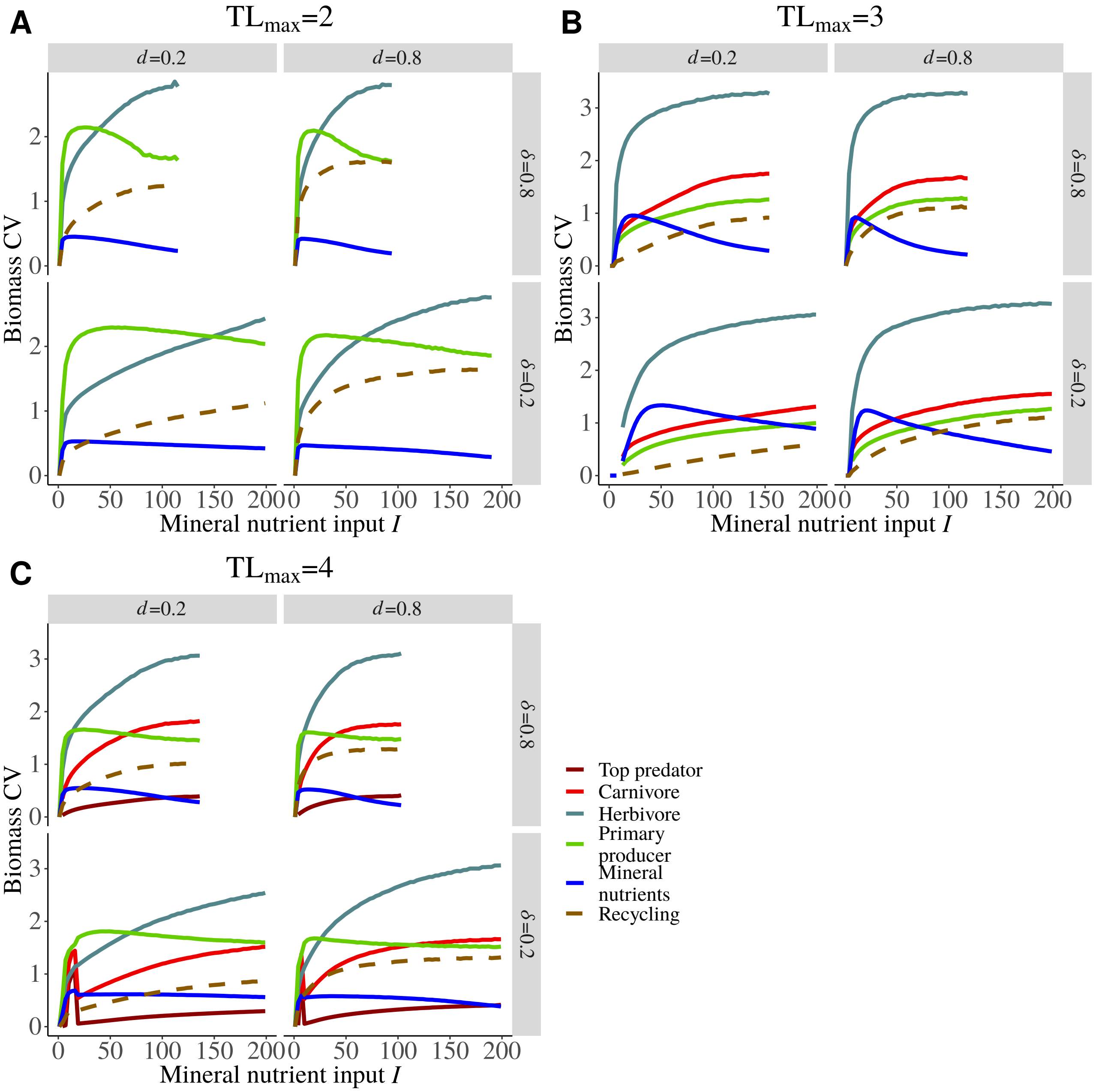


**Figure S2-10.** Biomass CV of recycled nutrients (dashed brown), mineral nutrients (blue), primary producers (green), herbivores (light blue), carnivores (red) and top predators (dark red) in C food chains. Three food chain lengths are tested: **A)** two species, **B)** three species and **C)** four species.

In addition, increasing food chain length increases or decreases species biomass time variability depending on their trophic level. For instance, primary producers are more variable when they are controlled by herbivores (Fig. S2-10A and S2-10C). This results are consistent with Shanafelt and Loreau (2018) who found that adding trophic levels to a food chain without nutrient cycling generates trophic cascades in both biomass and biomass CV. In fact, species at even distance from the top-consumer had a lower biomass and a higher biomass CV than when they were at odd distance from the top-consumer. However, our results are less clear with, for instance, the herbivore biomass CV that is always higher than carnivore biomass CV while Shanafelt and Loreau (2018) found that the herbivore biomass CV was alternatively higher or lower than the carnivore biomass CV with increasing food chain length. This suggests that nutrient cycling deeply changes species dynamics in food chains.

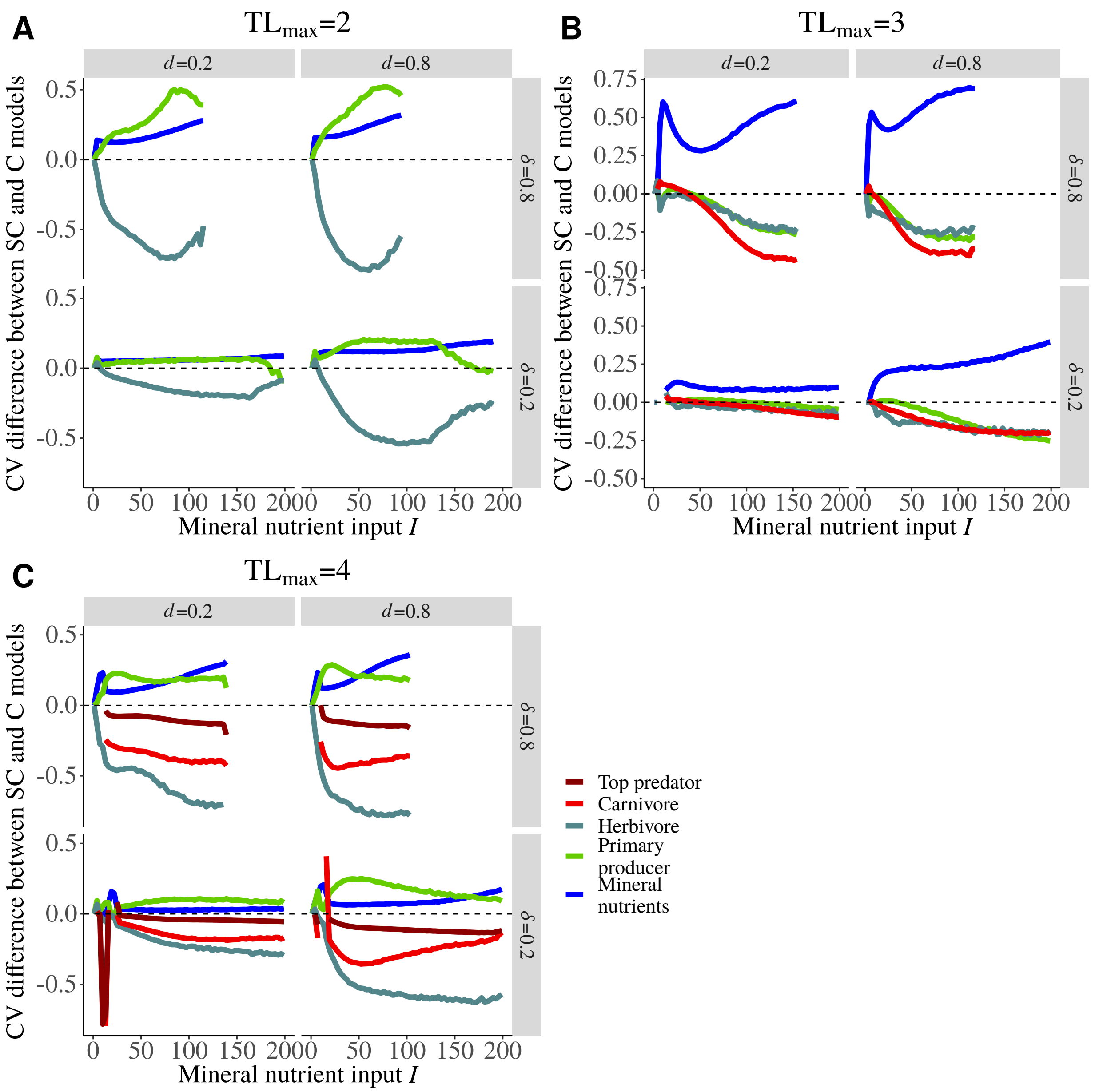


**Figure S2-11.** Difference between the biomass CV of mineral nutrients (blue), primary producers (green), herbivores (light blue), carnivores (red) and top predators (dark red) between SC food chains and C food chains. Positive values correspond to a higher CV in SC food chain and thus to a stabilising effect of nutrient cycling on dynamics. Three food chain lengths are tested: **A)** two species, **B)** three species and **C)** four species.

The presence of recycling feedback loops, once the enrichment effect of nutrient recycling is accounted for, has contrasting effects on species temporal variability depending on food chain length. Primary producers are stabilised for even food chain lengths (Fig. S2-11A and S2-11C), herbivores are destabilised whatever the food chain length, carnivores tend to be weakly stabilised at low nutrient input $I$ for $TL_{max}$=3 (Fig. S2-11B) and top predators are destabilised (Fig. S2-11C). More generally, primary producers have contrasting responses while consumers tend to be destabilised by nutrient cycling feedback loops. In addition, mineral nutrients are always stabilised by the presence of nutrient cycling feedback loops. As found in the food web model, the stabilising or destabilising effects of nutrient cycling feedback loops are stronger for high values of $d$ and $\delta$ and correspond to a higher quantity of recycled nutrients. This higher quantity of recycled nutrients (those directly recycled in particular) should intensify the coupling within the food chain and thus explain this increased effect on dynamics.

### S3 Appendix - Sensitivity analysis

#### Sensitivity to the way simulations were run

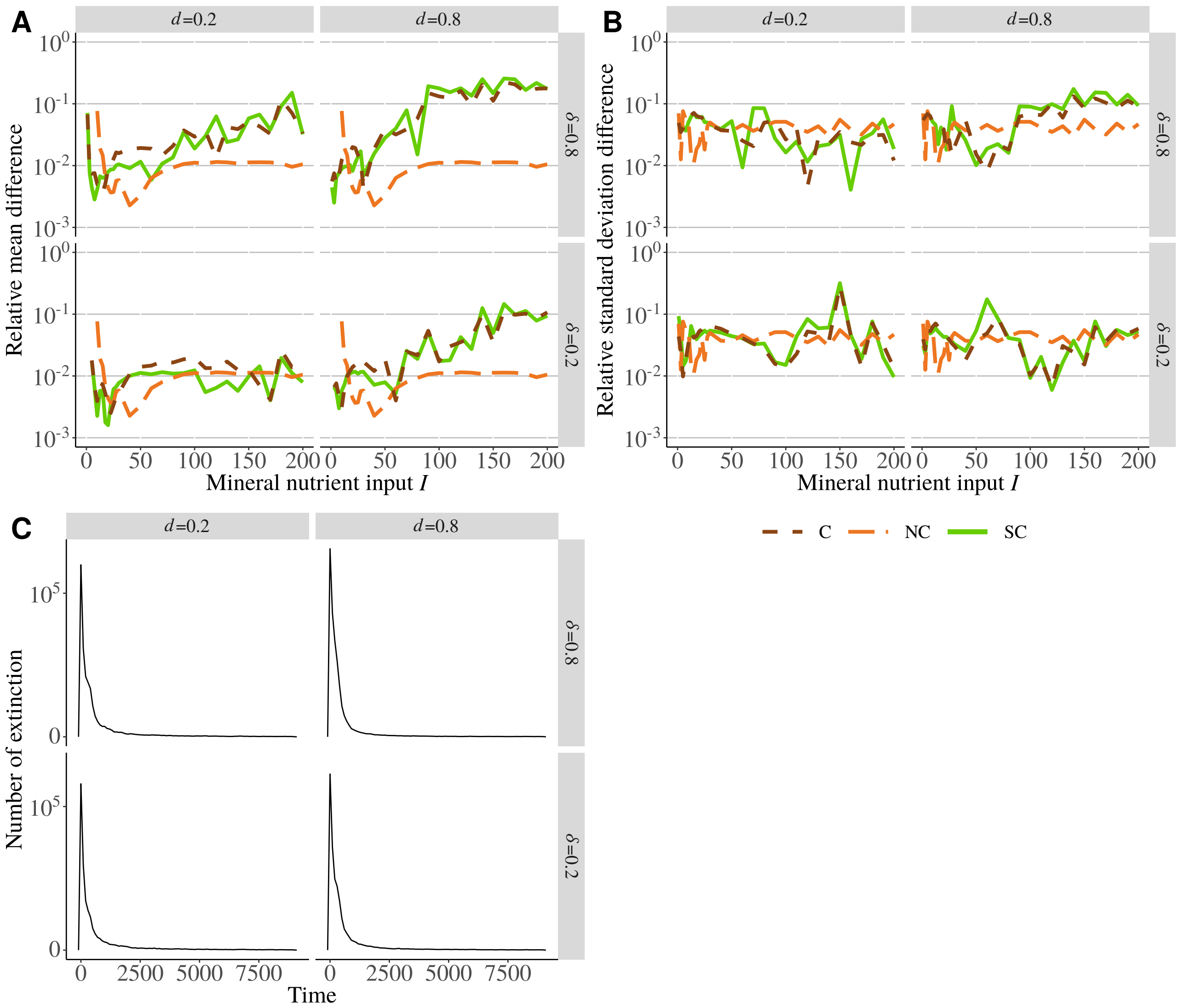


**Figure S3-1.** Sensitivity of the final community to the number of generated food webs and the number of time steps. **A)** Relative difference of the mean value of output variables (averaged for six variables: species persistence, average quantity of recycled nutrients, average primary and secondary productions, average quantity of nutrient directly recycled by primary producers and consumers) between simulations with 100 or 200 generated food webs for each combination of parameters. **B)** Relative difference of the corresponding standard deviations. **C)** Distribution of the number of extinctions over the 100 generated food webs along the transitory period. Extinctions are also cumulated over all the values of mineral nutrient input $I$.

As our food web model uses randomly generated communities, we average our different measures over 100 different communities (*i.e.* with randomly drawn body mass distributions). We assessed the number of required communities by calculating the relative difference between the means calculated with 100 or 200 different food webs for six variables. At low nutrient input we have a good precision for mean with a relative difference below 1% (Fig. S3-1A) and a difference lower than 10% for the standard deviation (Fig. S3-1B). The increase in the relative difference for the mean at high nutrient inputs must be due to the higher variability of food webs composition (*e.g.* maximum trophic level) due to multiple extinctions. Thus, 100 simulated food webs are enough to capture the accurate response of the model. Most extinctions occur within the first 2500 years of simulation, as shown by Fig. S3-1C. Thus, 9000 years are enough to get the final community and to get over the transitory regime.

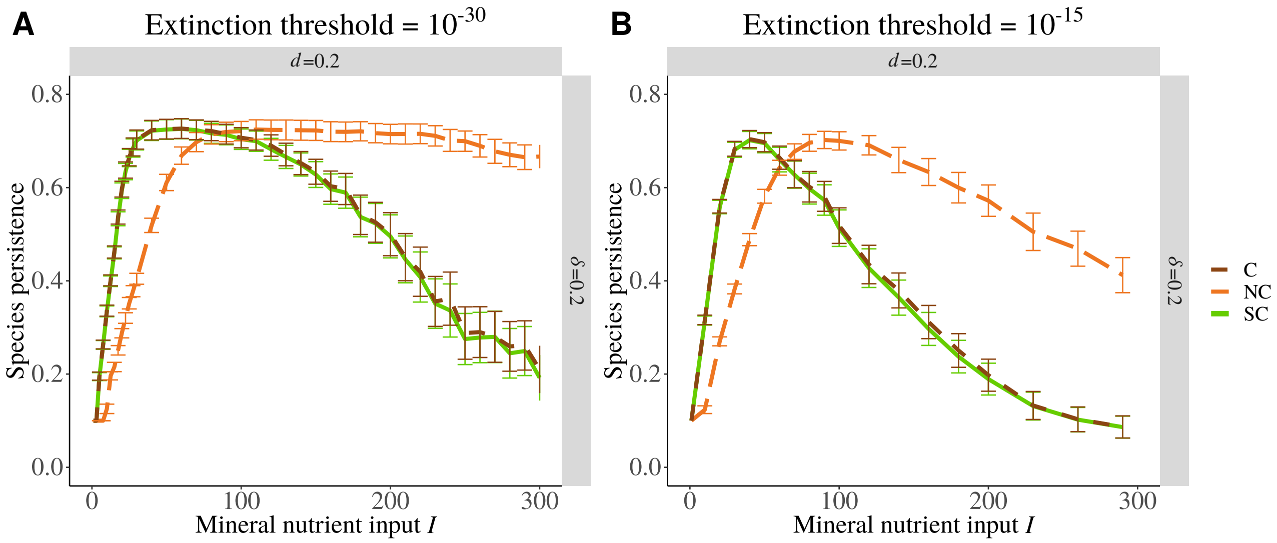


**Figure S3-2.** Sensitivity of the final community to the extinction threshold (*i.e.* the biomass under which the species is considered as extinct). **A)** Species persistence with an extinction threshold equal to $10^{-30}$ as in the main text. **B)** Species persistence with an extinction threshold equal to $10^{-15}$. 100 replicates are tested for each parameter combination.

We raised the extinction threshold up to $10^{-15}$kg.v$^{-1}$ (Fig. S3-2B) compared to the value used in the main study ($10^{-30}$kg.v$^{-1}$) (Fig. S3-2A). Species persistence is lower with this new threshold only at high nutrient inputs when species CVs increase with nutrient inputs. This demonstrates that extinction are due to an increase in oscillation amplitude that pushes species biomasses close to the extinction threshold.

#### Effects of attack rate and density dependent mortality rate allometric coefficients, of the nutrient loss rate and the half saturation of nutrient uptake

The decrease of species persistence with the decrease of the loss rate $\ell$ in Fig. S3-3A is due to the enrichment caused by the accumulation of nutrients. In fact, decreasing $\ell$ for a constant nutrient input $I$ increases the availability of mineral nutrients and is equivalent to an increase of nutrient inputs. The average CV of species biomasses (Fig. S3-3B) first increases with $\ell$ and then decreases. On the contrary, the half saturation of nutrient uptake $K$ only slightly affects species persistence and the CV of species biomasses when compared to $\ell$. For $\ell$ higher than $10^{-1.25}$ ($\sim 0.05$, corresponding to a loss of 5% of the nutrient stock), changing $\ell$ and $K$ does not affect species persistence and the CV of species biomasses. Then, we arbitrarily set $\ell$ and $K$ to maximise species persistence for $I \simeq 50$.

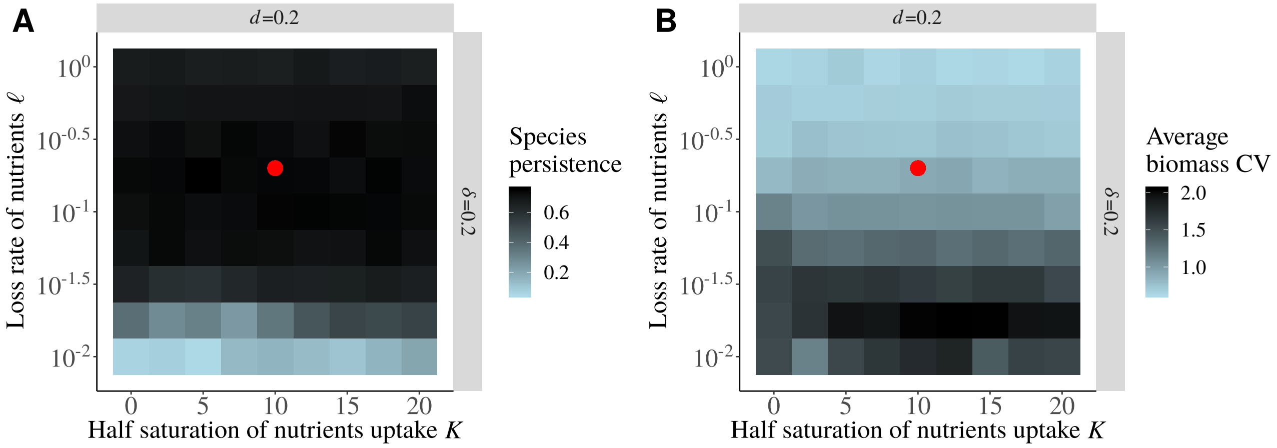


**Figure S3-3.** Effects of the half saturation constant of nutrient uptake $K$ and the loss rate of mineral nutrients and detritus $\ell$ on **A)** species persistence and **B)** the CV of species biomasses. Each square is the average of 100 simulated food webs (except for **B)** where only data from persistent food webs are represented). The mineral nutrient input is $I = 40$, the fraction of direct recycling is $\delta = 0.2$ and the decomposition rate of detritus $d = 0.2$. The red dots represent the combinations of parameters used in the main study ($K = 10$ and $\ell = 0.2$ in C and D).

We included a density dependent mortality rate ($\beta_i$) in our model to ensure a minimum species persistence. With a density dependent mortality rate allometric constant $\beta = 0$, the food web is so prone to the paradox of enrichment (see the higher average biomass CV in Fig. S3-4B) that no species can persist in the C model when recycling parameters are high (Fig. S3-4A). A high value of $\beta$ increases so much the death rate that strong nutrient inputs are needed to have a high species persistence (Fig. S3-4C) and biomass temporal variability is extremely low (Fig. S3-4D), thus resolving the paradox of enrichment. In addition, the average biomass CV is significantly higher in C models than in SC models for all combinations of $d$ and $\delta$, which differs to our results with intermediate values of $\beta$. However, the absolute value of biomass CV is so low ($10^{-5}$ that is even bellow the threshold used in Fig. 5 in the main text) that the effect on the overall dynamics seems negligible. Finally, whatever the value of $\beta$, the enrichment effect of nutrient cycling is always dominant to explain the difference between the C and the NC models as both curves representing the C and SC models overlap strongly as in Fig. 3B in the main text, making our results robust to $\beta$.

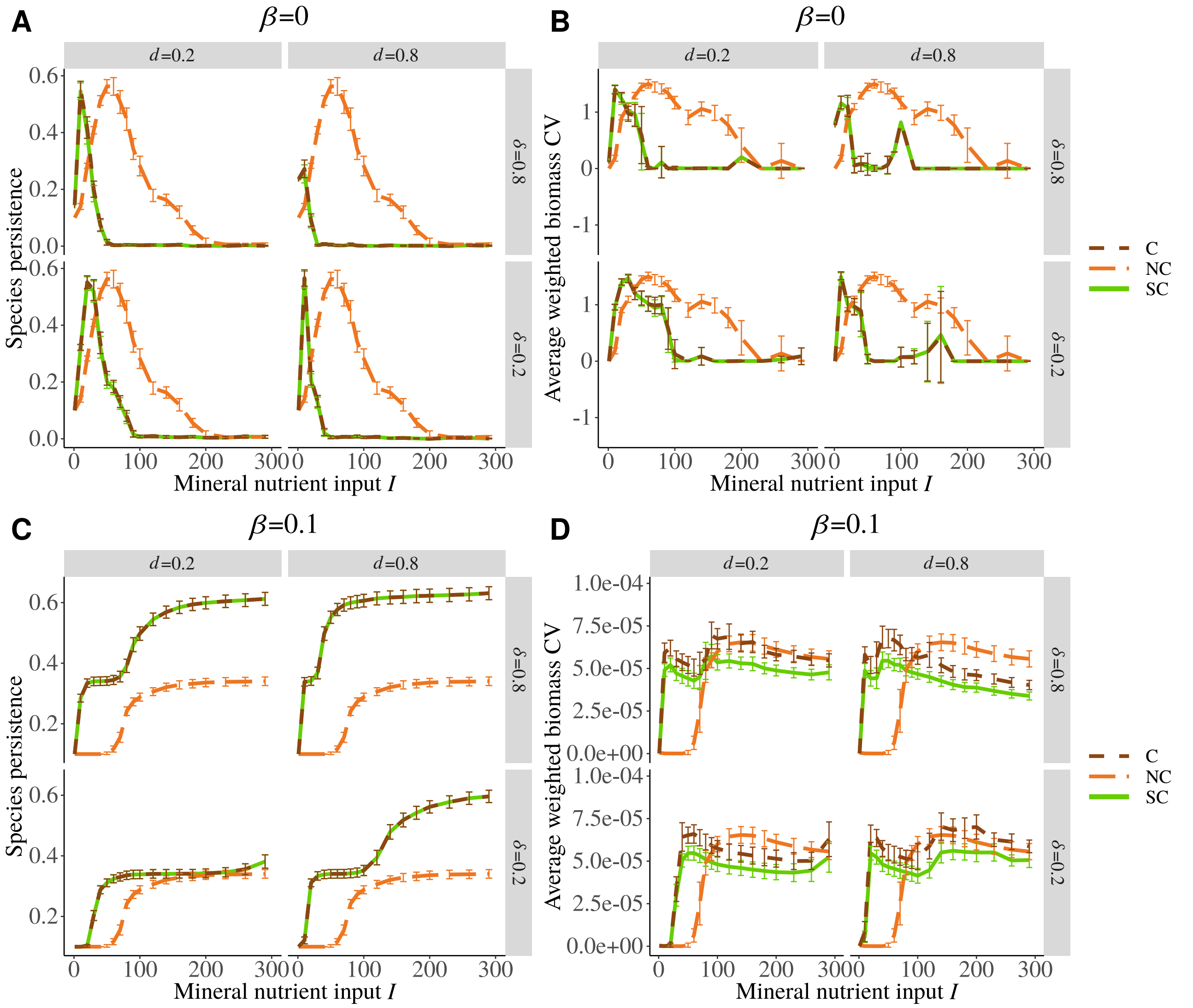


**Figure S3-4.** Sensitivity to the density dependent mortality rate allometric constant $\beta$. **A)** Species persistence and **B)** average weighted species biomass CV without density dependent mortality ($\beta = 0$) and **C),D)** with a strong density dependent mortality ($\beta = 0.1$). 100 replicates are tested for each parameter combination.

Species persistence in food webs is maximised only for restricted combinations of values of the attack rate allometric constant $a$ and the density dependent mortality rate allometric constant $\beta$ (Fig. S3-5A). In fact, if $a$ is high and $\beta$ low, consumers strongly exploit their prey and weakly self regulate, leading to the extinction of their prey and the collapse of the entire food web (Fig. S3-5F). The reverse combination leads to the extinction of consumers that cannot eat enough to compensate the loss of biomass due to a strong self regulation (Fig. S3-5E). The response of the CV of species biomasses to $a$ and $\beta$ is qualitatively similar to the response of species persistence (Fig. S3-5B) but occurs for smaller values of $\beta$. $\beta$ strongly dampens species biomass oscillations and a large part of the parameter space leads to food webs with very low average CV of species biomasses (regime similar to fixed points). Therefore, we chose $a$ and $\beta$ to maximise species persistence but with a minimal $\beta$. Indeed, a high $\beta$ strongly stabilise species dynamics and might obscure potential effects of nutrient cycling on stability.

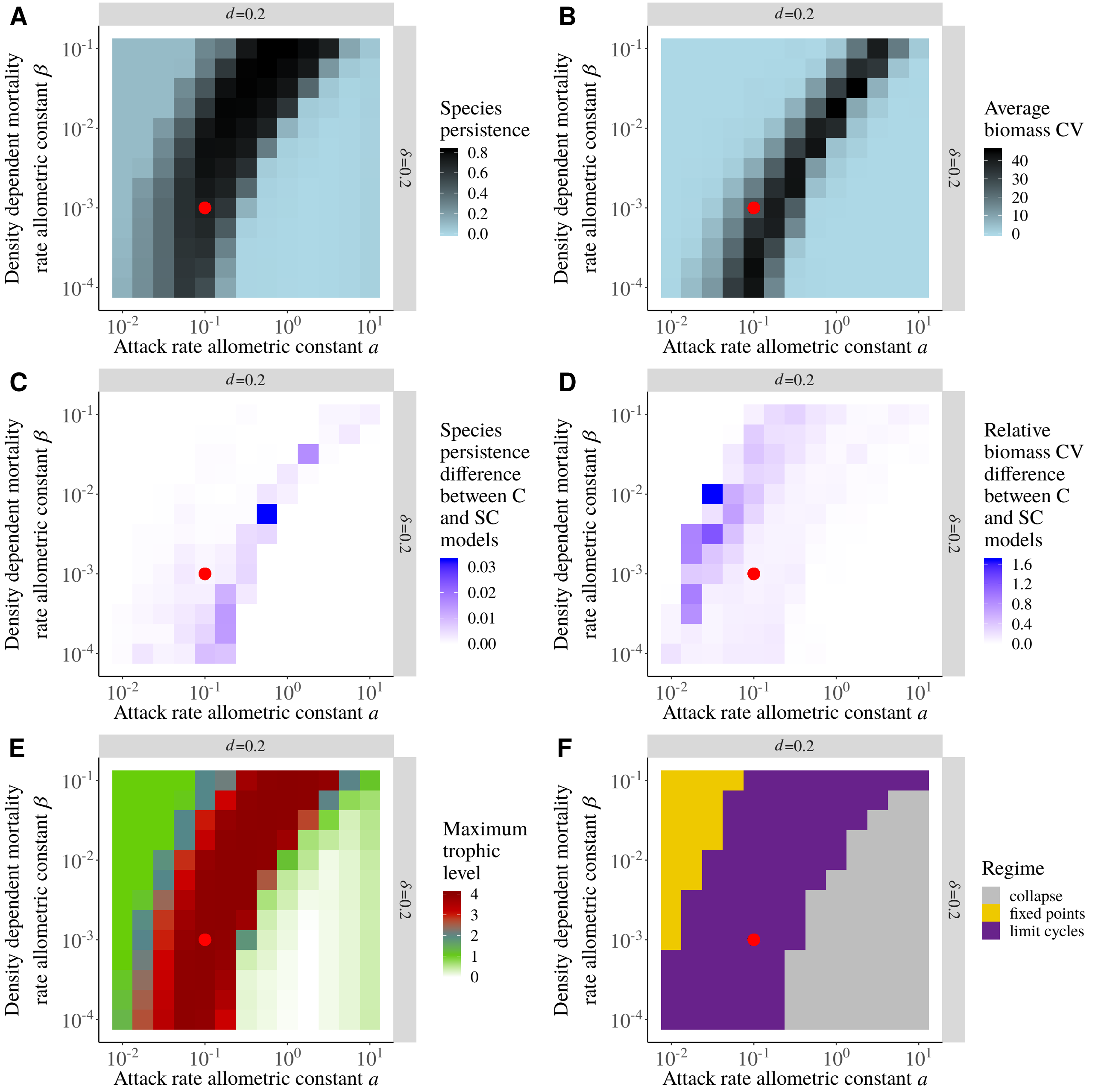


**Figure S3-5.** Effects of the attack rate allometric constant $a$ and the density dependent mortality rate allometric constant $\beta$ on **A)** the average species persistence and **B)** the average species biomass coefficient of variation. Average of the absolute value of the difference between the C and SC models for **C)** species persistence and **D)** species biomass CV. **E)** Average maximum trophic level. **F)** Regime of food webs that can display limit cycles or fixed points (average biomass CV lower than $10^{-4}$). When species persistence is mostly equal to zero, we consider that food webs collapse. Each square is the average of 100 simulated food webs (except for **B)**, **D)** and **E)** where only data from persistent food webs are represented). The mineral nutrient input is $I = 40$, the fraction of direct recycling is $\delta = 0.2$ and the decomposition rate of detritus $d = 0.2$. The red dots represent the combinations of parameters used in the main study ($a = 0.1$ and $\beta = 0.001$).

In spite of the high variability of species persistence and species biomass CV representing different possible regimes (fixed points or limit cycles in Fig. S3-5F) depending on the values of the attack rate allometric constant $a$ and the density dependent mortality rate allometric constant $\beta$, we do not see a significant difference between the responses of the C (with nutrient cycling) and the SC models (without nutrient cycling but with a simulated enrichment effect). Relative to the average values of species persistence (Fig. S3-5A) and biomass CV (Fig. S3-5B), the difference between the C and SC models are generally negligible (Fig. S3-5C and D). The significant differences occur only for species biomass CV when food webs are by the border between fixed point and limit cycle domains (Fig. S3-5F) as they can switch between food webs with only primary producers and food webs with consumers (Fig. S3-5E). Therefore, nutrient cycling mainly consists in an enrichment effect and weakly affects food web dynamics, whatever the value of the attack rate allometric constant $a$ and the density dependent mortality rate allometric constant $\beta$. Thus, our results are robust to the arbitrary choice of these parameters.

#### Sensitivity of the results to the value of adaptive foraging rate

We included adaptive foraging in our model as a mechanism promoting species persistence. Increasing the adaptive rate $A$ increases species persistence (Fig. S3-6A), as demonstrated by Kondoh (2003); Heckmann et al. (2012), while the qualitative response of species persistence to increased nutrient inputs remains unchanged (*i.e.* maximum of persistence occurring for the same values of nutrient input $I$). The CV of species biomasses (Fig. S3-6B) increases more sharply without adaptive foraging but the general response to nutrient enrichment remains qualitatively very similar for varying values of adaptive rate. In conclusion, our main results remain virtually unchanged when the rate of adaptive foraging is changed and we chose $A = 0.01$ as it promotes a high species persistence (higher values of adaptive rate only slightly increase species persistence).

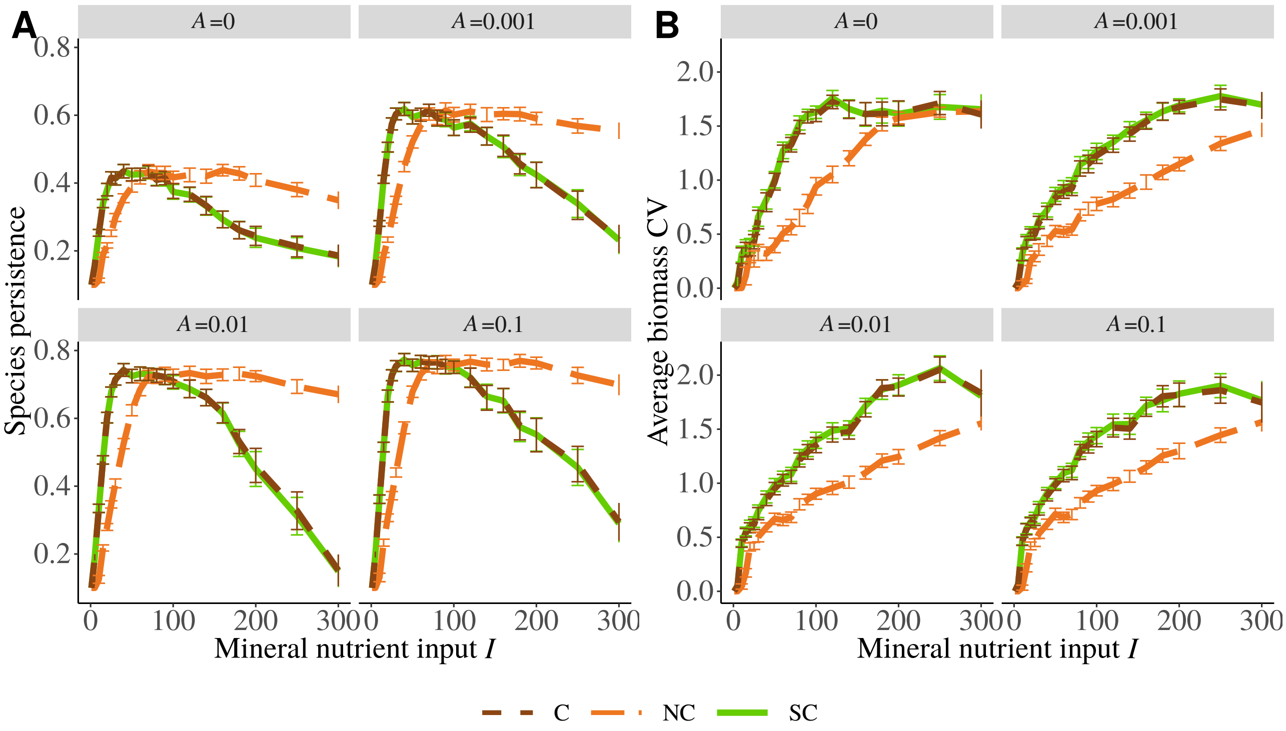


**Figure S3-6.** Effects of the adaptive rate $A$ of adaptive foraging on **A)** species persistence and on **B)** the average species biomass CV. We set $d = 0.2$ and $\delta = 0.2$, $A = 0.01$ is the value used in the main text. 100 replicates are tested for each parameter combination.

#### Sensitivity of the results to the type of functional response

The type II functional response leads to a decrease of species persistence at high nutrient inputs while species persistence stays maximum in food webs with a type III functional response (Fig. S3-7A). In addition, the CV of species biomasses is much lower in food webs with a type III functional response compared to food webs with a type II functional response. Thus, we do not observe a paradox of enrichment with a type III functional response in our model. This is consistent with the results of Rall et al. (2008). However, our results are qualitatively similar to those obtained with a type II functional response with a sharper increase of species persistence and a maximum of persistence reached for lower mineral nutrient inputs in C food webs than is NC food webs. We also still observe that the curves of the C and SC food webs strongly overlap. The major enrichment effect of nutrient cycling and its preponderance compared to the weak stabilising effect of positive feedback loops is thus robust to the type of functional response.

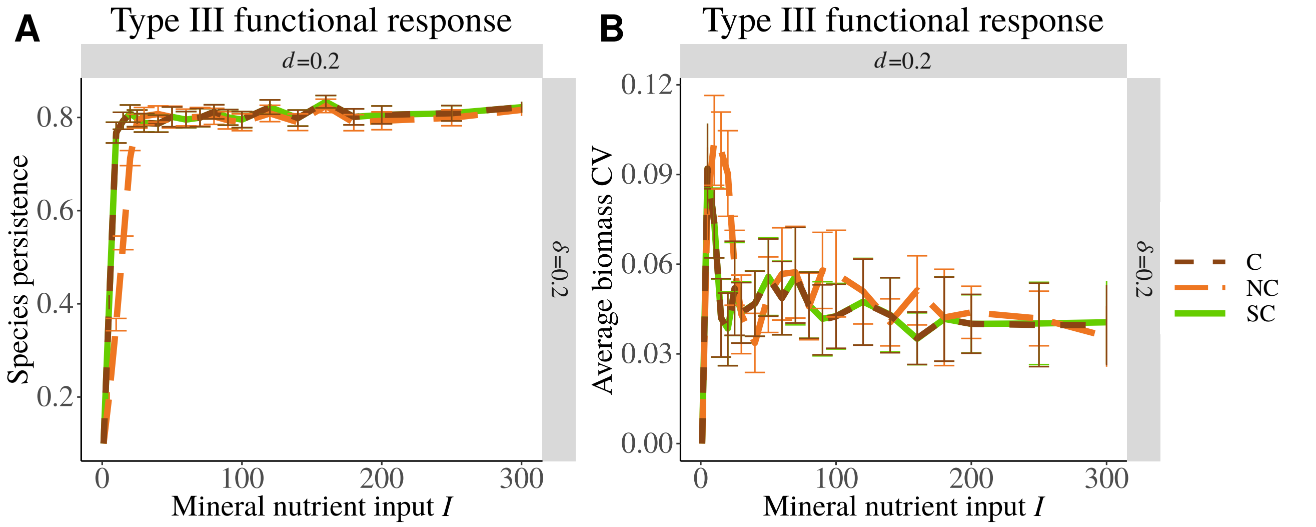


**Figure S3-7.** Effects of a type III functional response on **A)** species persistence and on **B)** the average species biomass CV. The type II functional response was used in the main text and 100 replicates are tested.

#### Sensitivity of the results to the C:N ratio of primary producers

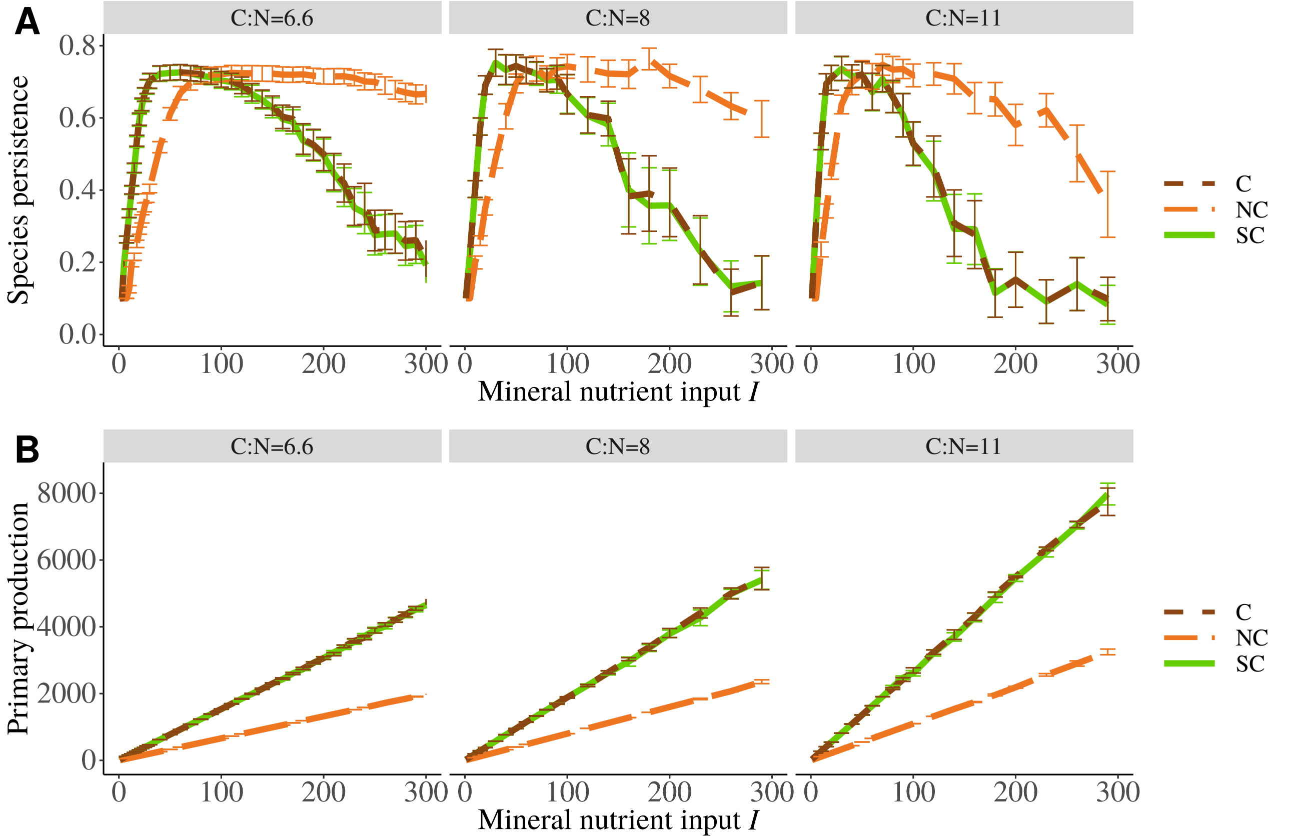


**Figure S3-8.** Effects of primary producers C:N ratio on **A)** species persistence and **B)** primary production. Primary producers C:N ratio is equal to 6.6 in the main text and 36 replicates are tested for C:N=8 and C:N=11.

The increase of species persistence (Fig. S3-8A) at low nutrient inputs and its decrease at high nutrient inputs are sharper if the C:N ratio of primary producers is high. The maximum species persistence is reached for lower nutrient inputs and the decrease of species persistence starts at lower nutrient inputs. This could be explained by the increase of primary production as the C:N ratio of primary producers increases (Fig. S3-8B). As the C:N ratio of primary producers increases, the growth of primary producers become less limited by the availability mineral nutrients. Therefore, increasing the C:N ratio of primary producers increases the productivity of the food webs and amplifies their response to nutrient enrichment.

In our model, the C:N ratio of primary producers does not strongly affects the response of the food web because the C:N ratio of detritus does not affect their decomposition. However, the C:N ratio of primary producers will be a central parameter in further models including a brown food web with decomposers (Attayde and Ripa, 2008; Zou et al., 2016) whose consumption rate strongly depends on detritus stoichiometry (Daufresne and Loreau, 2001).In nature, primary producer stoichiometry is highly variable between taxa but also within species depending on external conditions such as nutrient availability or light exposure (Sterner et al., 2002; Dickman et al., 2006; Danger et al., 2007, 2009; Mette et al., 2011). Its variations strongly impact ecosystem functioning through food quality and dead organic matter stoichiometry (Dickman et al., 2008; Cherif and Loreau, 2013).
